# Site-Dependent Decoupling of Drug-Biomarker Associations in Clear Cell Renal Cell Carcinoma Revealed by Functional Profiling of Patient-Derived Cell Models

**DOI:** 10.64898/2026.04.23.720088

**Authors:** Michaela Feodoroff, Tamara J. Luck, Romika Kumari, Minttu Polso, Patrick Penttilä, Minna Malmstedt, Piia Mikkonen, Lauren J. Gerber, Ruusu Merivirta, Mariliina Arjama, Erika Romppanen, Matilda Roos-Mattila, Pauliina Kallio, Swapnil Potdar, Mikaela Grönholm, Vincenzo Cerullo, Hanna Seppänen, Petrus Järvinen, Olli Kallioniemi, Tuomas Mirtti, Antti Rannikko, Vilja Pietiäinen

**Author notes:** Corresponding author: Vilja Pietiäinen. These authors share equal contribution (Michaela Feodoroff, Tamara J. Luck, Romika Kumari).

## Abstract

Clear cell renal cell carcinoma (ccRCC) frequently exhibits primary and acquired resistance to standard-of-care therapies, underscoring the need for improved strategies to predict therapeutic response and prioritize patient-specific treatments. Although recent multi-omic and single-cell studies have provided insight into the molecular landscape of ccRCC, molecular alterations alone incompletely predict drug sensitivity. We prospectively profiled tumors from 28 patients with localized and metastatic ccRCC by integrating molecular characterization, functional drug sensitivity and resistance testing in patient-derived cell models, and longitudinal clinical data. Although genomic biomarkers suggested potentially actionable therapies in 27/28 patients, functional testing revealed discordance between genomic actionability and ex vivo drug sensitivity, whereas linear mixed-effects modelling uncovered 16 novel copy-number-based features associated with sensitivity to 11 drugs. Genotype-drug response associations were largely preserved between primary tumors and vena cava thrombi but frequently disrupted in distant metastases. Integrating functional drug testing with multi-omic profiling reveals vulnerabilities not apparent from genomic data alone, refines therapeutic actionability, captures interpatient and intersite heterogeneity in ccRCC, and provides a scalable framework for individualized treatment prioritization.

## Introduction

Kidney cancer accounts for over 100,000 deaths worldwide each year^1,2^. Clear cell renal cell carcinoma (ccRCC), arising from the proximal renal tubule epithelium, represents approximately 85% of renal malignancies. Although surgical resection offers curative potential for localized disease, metastatic spread, which may occur synchronously or years after initial treatment, markedly reduces survival. Extension of tumor thrombi into the inferior VC reflects locally advanced disease, yet its biological and prognostic relationship to distant metastasis remains incompletely understood.

Genomically, ccRCC is characterized by loss of chromosome 3p and inactivation of the Von Hippel-Lindau (*VHL)* tumor suppressor gene, often accompanied by mutations in chromatin regulatory genes such as *PBRM1*, *BAP1*, and *SETD2*. These alterations drive profound transcriptional reprogramming^3^ but are rarely directly targetable. Consequently, current treatment strategies for ccRCC rely on surgical intervention and different pathway-directed, systemic therapies. Current systemic therapies including VEGF-targeted tyrosine kinase inhibitors (TKIs), immune checkpoint inhibitors (ICIs), and HIF-2α inhibitors have improved outcomes in advanced disease^4^; however, clinical responses remain heterogeneous, and predictive biomarkers to guide treatment selection are limited.

While recent multi-omic and single-cell atlases have provided detailed insights into the molecular landscape of ccRCC, translating these discoveries into individualized therapeutic strategies remains challenging. Molecular alterations alone often fail to fully predict treatment response, highlighting the need for complementary approaches that directly interrogate tumor vulnerabilities. Functional precision medicine, using patient-derived models to assess *ex vivo* drug sensitivity, offers a promising strategy to link genomic alterations with therapeutic response and guide personalized treatment^5,6^.

To address this gap, we established a prospective Development of Diagnostics and Treatment of Urological Cancers (DEDUCER) study for 28 patients with localized and metastatic ccRCC, including primary tumors, VC tumor thrombi, and distant metastases, integrating whole-exome sequencing (WES), and high-throughput drug sensitivity and resistance testing (DSRT) of patient-derived cell (PDC) models. By combining molecular and functional data, we identified both shared and patient-specific drug sensitivities, revealed discordance between genomic actionability and *ex vivo* drug response, and uncovered novel copy number-based associations with therapeutic vulnerability. Together, our findings demonstrate that functional drug profiling adds an orthogonal layer of information beyond genomic analysis alone and provides a scalable framework for refining therapeutic prioritization in ccRCC.

## Methods

### Study details, sample collection and clinical data

The study consisted of samples collected from 28 patients with ccRCC (22 males and 6 females), consenting for the ongoing prospective DEDUCER trial, registered at ClinicalTrials.Gov (NCT02994758). The study population comprised patients with ccRCC for whom clinical information was available (**Supplementary Table 1**): primary tumor and adjacent benign samples were collected at surgery from 23 patients. Beyond fresh primary ccRCC tissue, de novo VC tumor thrombi samples were obtained from seven patients and metastatic lesions (lung and small intestine) from two patients, all with matched primary samples. In addition, samples were obtained from a subcohort of five patients with pancreatic ccRCC oligometastasis as described earlier^7^. A board-certified pathologist verified histopathology for the obtained samples. All patients provided written informed consent, and the study was approved by the ethics committee of the Hospital District of Helsinki and Uusimaa (HUS) (Dnro 154/13/03/02/2016 (15.03.2017) and HUS/850/2017 (14.07.2021)). All patient material and clinical data were obtained as part of DEDUCER under Institutional Ethical Review Board-approved protocol and in accordance with the Declaration of Helsinki. The principles relating to the processing of personal data (GDPR) are considered in all data annotation and handling procedures.

### Cell culture

Cortical (PCS-400-011, ATCC) and proximal (PCS-400-010, ATCC) primary renal epithelial cells, together with ACHN (CRL-1611, ATCC), A-498 (ACC 55, DSMZ), Caki-1 (ACC 731, DSMZ), Caki-2 (ACC 54, DSMZ), and 786-O (CRL-1932, ATCC) cell lines, were cultured and maintained in accordance with the manufacturers’ instructions. Cell lines were authenticated by image-based phenotyping/Promega GenePrint24 System (Genomics Unit of Technology Centre, University of Helsinki, Finland) and tested negative for mycoplasma contamination using MycoAlert™ PLUS Mycoplasma Detection Kit (LT07-418, Lonza).

PDC cultures were generated from fresh cancer or adjacent benign tissue samples using the Tissue Dissociation Kit (Miltenyi Biotec). Cells were cultured in 2D with irradiated Swiss 3T3 mouse fibroblast feeder cells (J2 strain) (IR) on non-coated culture plates, as described earlier^8,9^ or on plates coated with 2% Matrigel (MG) (CLS354234, Corning) solution. 3D organotypic cell cultures were cultured in 0.2% or 0.4% GrowDex (GD) hydrogel (UPM Biochemicals) or 20% Growth Factor Reduced MG (CLS354230, Corning). The cultures were maintained in ROCK inhibitor-supplemented culture media (F-media or Basal media (BM)^9^ on standard or Primaria (PRIM) cell culture plates and dishes (Sarstedt #83.3920, #83.3902; Corning #353803) with or without MG coating and IR support.

### Immunofluorescence staining of PDCs

PDC cultures were stained with Ki-67 (Dako, #M724001-2), β-Catenin (Cell Signalling #8480) and carbonic anhydrase IX (CAIX, Abcam #10471), p53 (Dako #M7001) and phosphorylated mTOR (p-mTOR, Cell Signalling, #2974). Shortly, the PDCs were seeded into 384-well plates (Corning, #3764) and cultured for 2 days until subconfluent. Cells were fixed with 4% paraformaldehyde for 20 min at the room temperature (RT), washed with phosphate-buffered saline (PBS), and permeabilized with 0.3% Triton X-100 in PBS for 10 min at RT. Blocking was performed with 3% BSA in PBS for 1 h at the RT. Primary antibodies were diluted in 1% BSA/0.1% Tween-20 in PBS and added to the cells, followed by incubation for 1 h at 37 °C. After washing with PBS, secondary antibodies Alexa Fluor 488 anti-rabbit (Invitrogen, A11008) and Alexa Fluor 568 anti-mouse (Invitrogen, A11004) were applied at a final dilution of 1:2000 in 1% BSA/0.1% Tween-20 in PBS and incubated for 1 h at the RT. Nuclei were counterstained with Hoechst 33342 (Life Technologies, H1399; 1:10 000) for 5 min at RT. Washing steps were performed using a plate washer (Biotek, EL406). Cells were imaged using an Opera Phenix high-content confocal fluorescence microscope (Revvity; FIMM-High Content Imaging and Analysis (HCA) Core Unit, University of Helsinki) with a 20× water immersion objective (NA 1.0), acquiring nine fields per well.

### Exome sequencing

Genomic DNA samples of benign and cancer tissue, blood, and PDCs were processed in four separate experimental batches, using closely aligned workflows for library preparation, exome enrichment, and sequencing. Across the batches, between 50-150 ng of input DNA was used depending on extraction yield. For one batch involving cell suspensions, DNA was isolated using the Dynabeads® DNA DIRECT Universal kit (Thermo Fisher Scientific), and bead-bound DNA was used directly for library construction. All remaining batches used purified genomic DNA as input.

Library preparation was performed using either the KAPA HyperPlus kit (Roche) or Twist Bioscience library preparation chemistries, including the Twist Human Core Exome EF Multiplex Complete kit and the updated Twist Library Preparation EF 2.0 protocol with enzymatic fragmentation. Indexing adapters varied between batches and included Roche single index adapters, IDT unique dual index (UDI) adapters with UMIs, or PerkinElmer NEXTFLEX UDI adapters. Library quantification and quality control were performed using combinations of LabChip GX Touch HT High Sensitivity assays (PerkinElmer), Agilent Bioanalyzer High Sensitivity DNA assays, Qubit Broad Range DNA quantification (Thermo Fisher Scientific), and qPCR-based quantification kits (KAPA Library Quantification Kit or Thermo Fisher Collibri Kit). Libraries were pooled at 3-8 plex depending on concentration and batch.

Exome enrichment was performed using manufacturer’s recommended protocols with batch specific probe sets, including Roche MedExome probes, Twist Core Exome probes, Twist Human RefSeq spike in panels, and the Twist Exome 2.0 + Comprehensive Exome spike in set (37.45 Mb). All capture reactions followed Twist or Roche HyperCap 2.0 hybridization and wash conditions with minor workflow specific adjustments.

Sequencing was performed on Illumina platforms including HiSeq 2500 Rapid mode and NovaSeq 6000 systems using S2 or S4 flow cells depending on batch. Paired end read lengths were either 2 × 101 bp (standard format) or 101 + 10 + 10 + 101 bp for runs incorporating dual indexed 10 bp barcode reads under NovaSeq v1.5 chemistry. The sequencing was carried out at the FIMM Genomics NGS Sequencing Unit, University of Helsinki.

### Analysis of somatic and germline mutations and CNAs

Germline and somatic mutation analyses were performed on WES data using the Illumina DRAGEN Bio-IT Platform (versions v4.2.4, v4.0.3, and v3.9.3). Sequenced reads were aligned to the GRCh38 human reference genome. Prior to alignment, reads were trimmed to remove low-quality bases (minimum quality score of 3), poly-G tails, and TruSeq adapter sequences. Matched blood samples were used as the normal reference to distinguish somatic alterations. Somatic variants were identified using the somatic mode of DRAGEN’s small variant caller, which operates in tumor-normal mode and jointly analyzes tumor and matched normal samples. This approach assumes that germline variants and technical artifacts are shared between both samples, whereas true somatic mutations are unique to the tumor sample. Predicted somatic variants were functionally annotated using ANNOVAR^10^.

Gene copy number alterations (CNAs) were identified using the CNV kit^11^ by comparing tumor/PDC samples against matched normal (blood) samples. Read depth was calculated for target and anti-target regions, normalized using a pooled normal reference, and corrected for systematic biases. Segmentation was performed using circular binary segmentation (Implemented via DNA copy) to identify regions of CNAs. Further, the obtained Log2 CN ratios across the genomic bins were mapped to the corresponding genomic cytobands using bedtoolsr package^12^. Segments with absolute log2 CN ratios ≥ 0.2 were classified as gains or losses for downstream analysis. Loss of heterozygosity (LOH) events and allele-specific CN calls were identified using ASCAT, using default parameters and tumor/normal paired samples^13^.

### Mutation frequency comparison with TCGA-KIRC reference cohort

Somatic mutation frequencies in the primary tumor, VC tumor thrombi and metastatic samples were each compared to frequencies in the TCGA-KIRC primary ccRCC cohort (n=451), using the “Kidney Renal Clear Cell Carcinoma (TCGA, Firehose Legacy)” dataset downloaded from cBioPortal^14,15^. For each gene and sample group, Fisher’s exact test (FET) was used to compare frequencies. False discovery rate was used to adjust for multiple testing across genes.

### Clinical actionability analysis using PCGR

The PASS-filtered DRAGEN VCFs (for somatic mutations) and ASCAT segmentation files (for somatic CNAs) were analyzed using the Personal Cancer Genome Reporter (PCGR) (version 2.2.5; tumor primary site “kidney”, data bundle 20250314)^16^. PCGR results were used to i) classify somatic variants based on their clinical actionability using the AMP/ASCO/CAP framework, and ii) determine underlying mutational signatures, tumor mutation burden (TMB) and microsatellite instability (MSI) status.

### WES-based estimation of tissue and PDC tumor cell content

ASCAT^13^ was used with default parameters and tumor/normal paired sample analysis to obtain WES-based estimates of tumor purity/tumor cell content. Subsequent manual review of ASCAT sunrise plots identified samples with tumor purity estimates of low confidence, i.e. lack of distinct local optimum on the tumor purity y-scale. For samples with high confidence findings, ASCAT (CNA-based) tumor purity estimates were used ’as is‘. For samples with low confidence ASCAT purity calls, the maximum variant allele frequency (VAF) of each sample’s key somatic driver mutations was used instead (i.e. variants classified as PCGR Actionability Tier1-2^16^ and/or coding variants in known ccRCC driver genes *VHL, KDM5C, PBRM1, SETD2, BAP1*, and *TP53*).

### Drug sensitivity and resistance testing

DSRT was performed on PDCs from 14/28 patients (P1, P2, P4, P7, P10, P12, P13, P14, P15, P16, P22, P24, P25 and P29) derived from primary tumor, benign adjacent tissue, VC tumor thrombi and metastatic tissues. Primary renal cortical and proximal tubular epithelial cells derived from non-malignant tissue were included as non-tumor reference models of normal kidney epithelium, together with commercial renal cancer cell lines (ACHN, A498, Caki-1, Caki-2, and 786-O). DSRT experiments were conducted using low-passage cultures (passages 1-8) across different culture conditions (**Supplementary Table 2**). A collection of up to 535 approved and investigational compounds (**Supplementary Table 3)**, were plated on 384-well plates using Echo Acoustic dispenser in five different concentrations in a clinically achievable range and diluted in 0.1% dimethyl sulfoxide (DMSO) or water. DMSO (0.1%) and benzethonium chloride (100 µM) served as negative, and positive controls, respectively. Cells were seeded in single cell suspension (500-2000 cells/well) on pre-drugged flat bottom (2D-DSRT, Corning #3674) or U-bottom (3D-DSRT, Corning #3830) 384-well TC plates with robotic handling (Agilent BioTek MultiFlo FX BRAD dispenser). Cell seeding was followed by centrifugation (300 x g, 20 s) to remove any air bubbles. Plates were then incubated for 72 h in a humidified 5 % CO2 incubator at 37°C. Prior to the experimental endpoint with cell viability measurement, CellTiter-Glo reagent (Promega) was dispensed (Agilent BioTek MultiFlo FX BRAD dispenser) at 15-25 µL volume to each well. Plates were shaken on an orbital plate shaker for 5 min, centrifuged for 5 min and incubated for 20 min at the RT, as described previously^17^. The data was run through the quality analysis pipeline to confirm the robust Z prime (>0.70) for the assay (in-house analytics tool Breeze)^18^, and DSS were calculated for each compound to quantify drug efficacy across the tested concentration range^19^.

For downstream heatmap analysis to identify drug response patterns across samples, only representative PDC models were included. DSRT replicate measurements were averaged (mean) for each matching WES sample, including replicates screened under both 2D and 3D culture conditions using the FS2A (116-drug) and FO5A (528-drug) screening plates. Drugs with a mean DSS > 10 in at least one sample and with fewer than 5 missing values (NAs) across all samples were retained for analysis. To assess intra-sample-type heterogeneity in drug responses, DSS were compared across patients within each biological sample category, including tumor (T), vena cava thrombus (VC), metastasis (MET), benign samples (B), primary cortical/proximal cell cultures, and cancer cell lines. Drug response distributions were visualized to evaluate variability in sensitivity patterns within and between sample groups. Statistical differences in DSS distributions among patients within each sample-type category were assessed using non-parametric Kruskal-Wallis tests. For the pairwise comparison of DSS between matched 2D and 3D DSRT for each individual patient sample (n = 11), drugs with fewer than 5 missing values (NAs) across all samples, regardless of the mean DSS, were retained for the analysis. For each of these patients, DSS values obtained under 2D and 3D culture conditions in DSRT were compared using a paired Wilcoxon signed-rank test.

Differential drug sensitivity analyses were performed between T, VC, and MET samples using limma framework. A no-intercept design matrix was specified to model group-specific means for T, VC, and MET samples (∼0+Group). To account for non-independence of multiple samples from the same patient, patient identity was included as a blocking factor, and within-patient correlation was estimated using duplicateCorrelation. Linear models were then fitted with this correlation structure (lmFit, with block and correlation), and predefined contrasts were tested (T vs VC, T vs MET, and VC vs MET). Contrast-specific statistics were obtained using contrast.fit followed by empirical Bayes moderation (eBayes). Drugs with an adjusted p-value < 0.05 and an absolute delta DSS > 5 were considered significantly differentially sensitive.

To investigate drug response patterns at the pathway and target-class level, drugs were grouped according to their annotated functional classes. DSS were compared across functional classes and sample categories. DSS distributions within each functional class were visualized to evaluate variability in pathway-level drug sensitivities across sample types. Functional classes were ranked according to median DSS values within each sample category. To further characterize group-specific drug response patterns, median DSS values were calculated for each drug across sample type. Heatmaps based on median DSS values were generated to identify functional drug classes and compounds enriched in specific sample groups.

### Novel biomarker identification

R packages lme4 (version 1.1.35.5^20^) and lmerTest (version 3.1.3^21^) were used to generate linear mixed-effects models (LMMs) and test our validated PDC models for significant associations between individual genes’ somatic VAFs or copy number ratios (CNR) and individual drugs’ DSS values, while accounting for tissue type (benign/tumor/VC/metastasis) and patient effects. First, drugs and genomic alterations were filtered to reduce the multiple testing burden: Drugs were restricted to those with selective DSS (sDSS, defined as PDC DSS - mean Primary Cortical/Proximal cell DSS) ≥ 10 in at least one sample, to focus on drugs effectively killing PDCs rather than non-cancer control cells *ex vivo* (40/535 drugs retained). Somatic gene mutations were restricted to those affecting protein sequence (coding splice-site/exonic non-synonymous variants), mutated in at least two patients, with VAF ≥ 10% in at least three samples and affecting genes in the COSMIC cancer gene census (harboring 764 cancer-relevant genes^22^) (38 genes retained). Somatic CN-altered genes were restricted to coding genes (excluding genes located on chrX and chrY), harboring CNAs (mean log2 CNR > log2(1.25) | mean log2 CNR < log2(0.75)) in at least three samples and two patients and part of the COSMIC cancer gene census (400 genes retained). Using a final model of ‘DSS_drugA ∼ VAF_geneB + TissueType + (1 | PatientID)’ for each drug/somatic mutation pair (40 drugs * 38 genes = 1,520 models total) and ‘DSS_drugA ∼ log2CNR_geneC + TissueType + (1 | PatientID)’ for each drug/somatic CNA pair (40 drugs * 400 genes = 16,000 models), p-values were calculated for each gene/drug pair association and adjusted for multiple testing. Of the 17,520 total models, 56 retained significance at FDR <0.1. To ensure that these 56 associations were robust and not driven by individual patient outliers, we performed Leave-One-Patient-Out (LOPO) cross-validation. Specifically, significant drug/gene pairs were each re-analyzed 14 times, each time withholding all the samples from one of the 14 patients for which DSRT data was available. Finally, 19/56 associations were deemed “robust”, as they maintained a consistent direction of effect and remained statistically significant (p < 0.1) across all LOPO iterations.

### Biomarker validation using DepMap

For 16 of 19 robustly significant drug/gene pairs with respective available data, associations were validated using *ex vivo* drug response and copy number data of six primary ccRCC-derived cancer cell lines from DepMap (KMRC1, KMRC3, KMRC20, 786O, OSRC2, 769P)^23,24^. Specifically, the PRISM Repurposing Public 24Q2 DSRT data^25^ was used. As DepMap drug response data was quantified using log2FC cell viability of drug-treated vs DMSO-treated controls, where lower values are associated with increased drug sensitivity, whereas our DSRT data was quantified using DSS values, where higher values indicate increased drug sensitivity, DepMap data was converted to -logFC growth inhibition, to make trend directions more directly comparable (higher value of both scales indicating higher drug sensitivity).

For all 16 distinct genes among the 19 drug/gene biomarker pairs, associations between gene copy number and mRNA expression were investigated using the same six ccRCC-derived cancer cell lines from DepMap. Specifically, the DepMap Public 25Q3 dataset of log-transformed transcripts per million (TPM) of protein-coding genes was used. First, regression coefficients (slopes) of association were determined for each gene using linear regression (R’s lm() function), with “positive associations” defined as slopes >0. To assess enrichment of positive associations among the 16 biomarker genes, an empirical p-value was calculated using 1000 random gene set permutations: For each permutation, a random set of 16 genes was sampled from the 18399 total genes for which copy number and mRNA TPM data were available in DepMap, and the number of genes with positive TPM∼CNR slopes was determined. Among the 1000 permutations, we found 7 with at least 15/16 genes having positive slopes, i.e. p=0.007.

### Drug target-biomarker network analysis

To investigate the mechanistic relationship between the predicted biomarkers and drug action, we reconstructed signaling subnetworks connecting direct drug targets to the corresponding candidate biomarkers using the OmniPath prior-knowledge interaction network^26^. Transcriptional and post-translational interactions were retrieved from the OmniPath webserver and shortest path subgraphs were assembled using the igraph R package (version 2.3.2) for each biomarker-associated drug by connecting its molecular targets to the corresponding predicted biomarkers. Direct drug targets were compiled from in-house annotations based on DrugBank^27^, TTD^28^ and ChEMBL^29^.

### Statistics

Statistical analyses were performed using R version 4.5.1 (R Core Team, 2025)^30^.

## Results

### Sample collection and characteristics of ccRCC patient cohort

We established an integrated translational framework in 28 patients with primary/metastatic ccRCC, combining longitudinal clinical annotation with matched biospecimens and patient-derived functional models (**Figure 1**; for clinical information see **Supplementary Table 1**). Surgically resected specimens were processed immediately to generate a living biobank, enabling parallel histopathological and genomic analyses. In addition, enzymatically dissociated tissues were further used to establish low-passage *in vitro* culture models for functional interrogation of patient-specific tumor biology.

**Figure 1.**
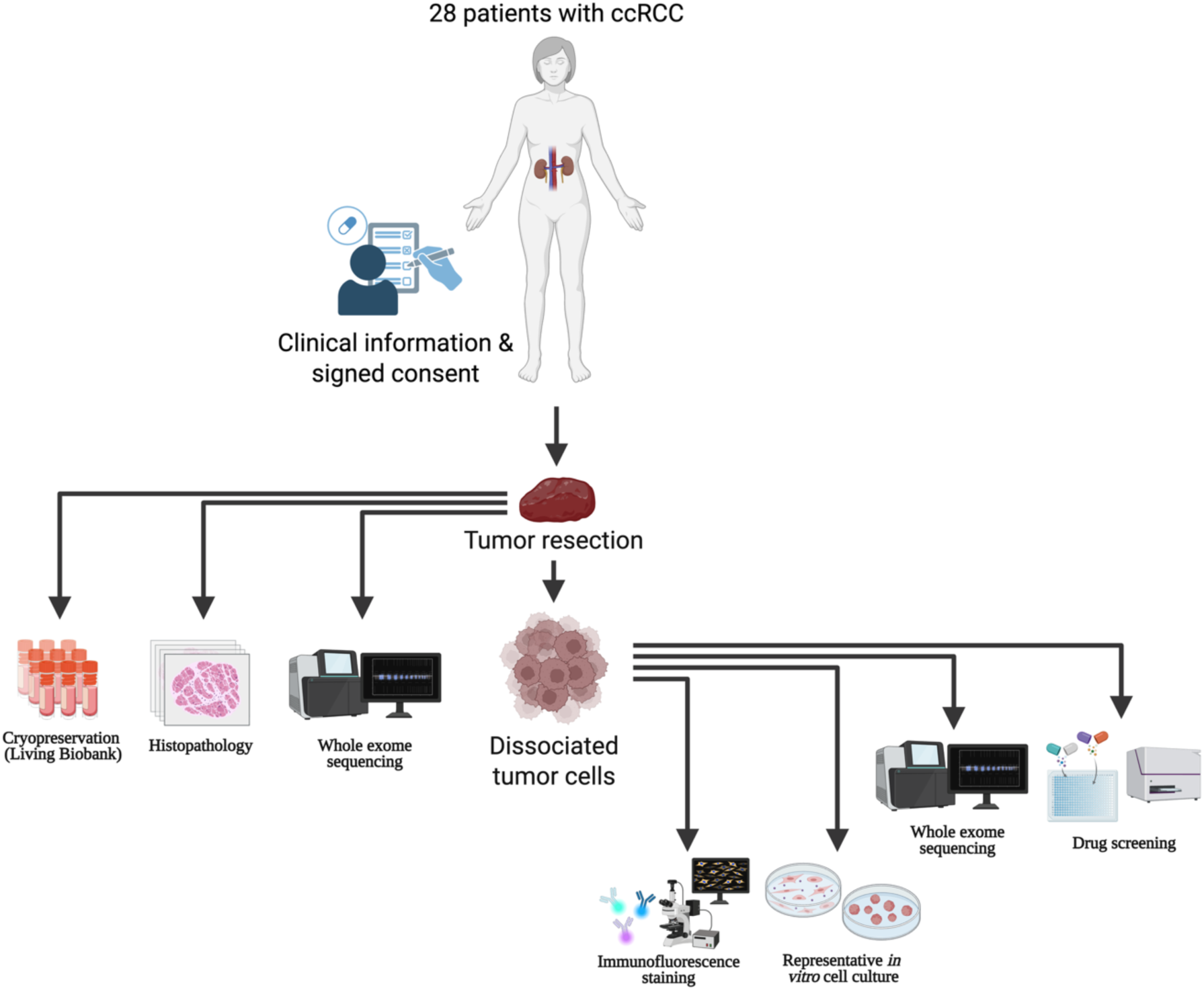
Functional precision medicine study overview. Schematic of the clinical and biospecimen data collected from 28 patients with ccRCC. Freshly resected tissues were processed to establish a living biobank and subjected to histopathological evaluation and WES. Dissociated tissues were used to generate patient-derived cell models for downstream characterization, including immunofluorescence staining, WES, and drug sensitivity and resistance testing. Created with BioRender.com

These models were further allocated to downstream experimental pipelines, including immunofluorescence-based phenotypic characterization, WES, and DSRT. The integrated framework enabled direct linkage of clinical features, molecular alterations, and functional drug response data within individual patients. The framework provided a flexible platform for multi-layered investigation of tumor biology in a clinically annotated prospective cohort.

### Recurrent ccRCC driver mutations are preserved across primary, VC, and metastatic tissues

WES was performed on 38 surgically resected fresh tissue samples from 28 patients in our ccRCC cohort. *VHL* mutations were detected in 18 of 23 primary tumors (78%). *VHL* alterations were also identified in 4 of 7 VC tumor thrombi (57%) and 6 of 7 metastatic samples (86%), typically in the same patients who had VHL mutations also in the primary tumor (**Figure 2A; Supplementary Figure 1A)**. These somatic *VHL* variants comprised, most commonly, missense (n=12), frameshift (n=11), and stop-gain (n=4) mutations. Beyond *VHL*, recurrent mutations were observed in additional ccRCC-associated genes. *PBRM1* mutations were present in 10 primary tumors (43%), 3 VC tumor thrombi samples (43%), and 4 metastatic samples (57%), while *SETD2* mutations were identified in 6 primary tumors (26%), 2 VC tumor thrombi (29%), and 5 metastatic samples (71%). These frequencies are consistent with previously reported genomic landscapes of ccRCC^31^.

**Figure 2.**
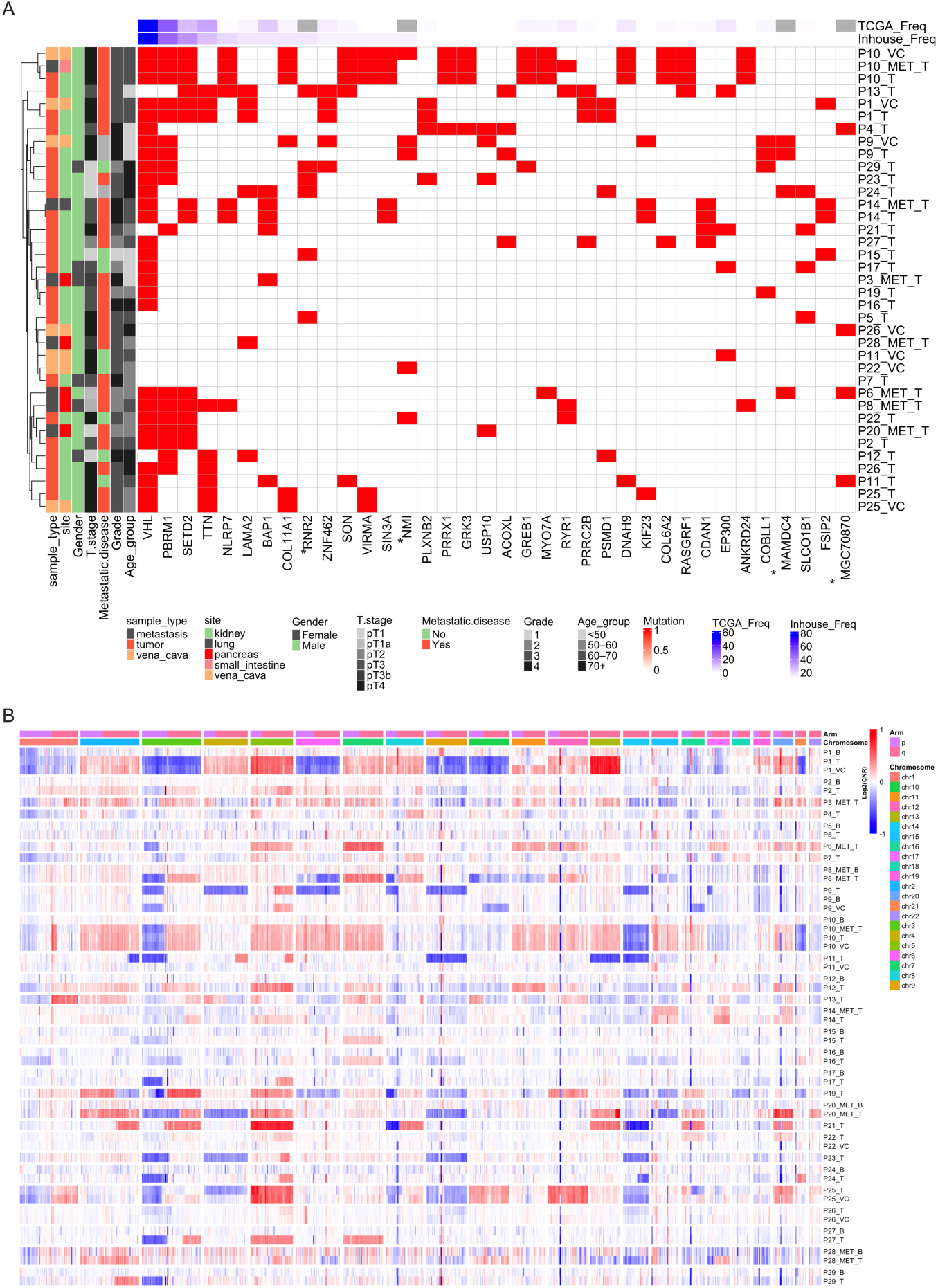
Genomic landscape of the cohort highlighting somatic coding mutations and copy number alterations. **A.** Protein-coding gene mutations observed in >3 tumor samples, across 28 patients (38 samples). Genes are ordered by mutation frequency of each gene. The matrix indicates mutation presence (1) or absence (0) per sample. Genes labeled with * were not found to be mutated in the TCGA ccRCC dataset. **B.** Chromosome-level CNA summary across the genome for primary tumor (T), metastatic (MET), adjacent benign (B), and VC tissue samples. Heatmap shows log2(CNR), where red indicates copy number gain, blue indicates copy number loss, and white indicates neutral copy number.

Overall, recognized driver variants were rarely gained in VC tumor thrombi or metastatic samples relative to the matched primary tumor. Among the 7 VC tumor thrombi and 7 metastatic samples sequenced, we only observed one VC and one metastatic sample each with an additional driver variant not detected in the respective primary tumor (additional *TERT* variant in P14 metastasis; additional *TSC1* variant in P26 VC tumor thrombi) (**Supplementary Figure 1A**).

To further contextualize our cohort, we compared somatic mutation frequencies of genes recurrently mutated in >3 patients in our cohort (**Figure 2A**) across primary tumors, VC tumor thrombi and metastatic samples. Primary tumors were enriched for *ACOXL* (n=4/23), *PSMD1* (3/23) and *SLCO1B1* (4/23) mutations (compared to 1-3 mutated cases out of 451 in TCGA-KIRC; Fisher’s exact test, FDR = 0.001-0.004). Metastatic samples showed enrichment of *SETD2* (5/7 vs 51/451, FDR = 0.006) as well as *NLRP7*, *SIN3A* and *ANKRD24* (2-3/7 vs 1-3/451, FDR = 0.001-0.009) (**Supplementary Table 4**). No enrichment was found for these recurrently mutated genes among our VC samples compared with TCGA-KIRC. In addition, our cohort exhibited recurrent mutations in several genes that were not reported for the TCGA cohort, including RNR2, NMI, MAMDC4, and MGC70870 (**Figure 2A**). Collectively, these findings confirm that canonical ccRCC driver mutations are preserved across primary tumors, VC tumor thrombi, and metastatic lesions in our cohort, while highlighting selected site-specific enrichments.

### Tumor mutational burden is stable across anatomical ccRCC tissue sites

TMB was assessed across the cohort. Most samples exhibited low to moderate mutational loads, with median values of 3-4 somatic mutations/Mb across primary tumors, VC tumor thrombi, and metastatic samples, compared with a median of 0.15 mutations/Mb across the benign samples (**Supplementary Figure 1B**). While TMB was significantly lower in benign compared with tumor samples (Wilcoxon p = 2.8x10^-7^), no significant differences were observed between primary tumors and VC tumor thrombi (p = 0.57) or metastatic tissues (p = 0.51). Interestingly, one primary tumor (P13_T) demonstrated a markedly elevated TMB (22.5 mutations/Mb). Despite this, the sample harbored canonical ccRCC alterations, including a *SETD2* mutation (**Figure 2A**) and loss of chromosome 3p (**Figure 2B**). In addition, P13_T displayed a distinct mutational signature associated with Aristolochic acid (AA) exposure, alongside the more typical ccRCC signatures observed across the rest of the cohort (**Supplementary Figure 2**). On the basis of the characteristic ccRCC mutations and copy number variations, the sample was not excluded from further analysis.

### Clonal heterogeneity and landscape across ccRCC tissues

We next assessed clonal heterogeneity by examining VAFs of clinically relevant somatic driver mutations across primary tumors, VC tumor thrombi, and metastatic tissues. Analyses focused on clinically actionable Tier 1-2 variants (as defined by PCGR)^16^ and mutations in canonical ccRCC drivers *VHL*, *PBRM1*, *SETD2*, *DNAH9*, *KDM5C,* and *BAP1* (**Supplementary Figure 1A**).

Marked inter-individual variability was detected across the cohort. Several patients (n=9/28) exhibited high driver VAFs (≥0.5), consistent with dominant clonal mutations, whereas others (n=14/28) showed exclusively lower VAFs (<0.3), even for canonical drivers such as *VHL*, suggesting subclonality or reduced tumor purity. WES-based computational estimates of sample tumor cell content correlated well with overall VAF patterns of these clinically relevant driver VAFs (gray shading in **Supplementary Figure 1A**). The number of driver mutations per patient also varied substantially, ranging from one or two events to multiple concurrent alterations. Intrapatient comparisons revealed that in five of the eight cases for which multiple cancer tissues had been sequenced (P1, P9, P10, P14, P25), VAFs were largely conserved across anatomical sites, indicating stable clonal representation between primary tumors, VC tumor thrombi, and metastases. In contrast, the other three patients demonstrated limited overlap in mutation VAFs across tissues, especially in the presence of lower tumor cell content of either tissue sample (P11, P22, P26).

### Recurrent CNAs are preserved across primary, VC, and metastatic tissues

Copy number analysis (CNA) revealed widespread genomic alterations across ccRCC tissues (**Figure 2B**). The most frequent gains (>25% of samples) involved chromosome 5, including 5q (42.5%) and 5p (40%). Additional recurrent gains were observed on 2q (25%), and 20p (25%). Among recurrent losses (>25% of samples), chromosome 3p (50%), harboring the VHL gene, was the most prominent. Deletions of 21p (40%), 22p (32.5%), 14q (32.5%), and 9q (25%) further defined the characteristic CNA profile of this cohort. Overall, across the cohort, we observed extensive large-scale CNAs, with largely conserved patterns among tumor, VC, and metastatic samples from the same patient, whereas matched benign samples remained largely copy number neutral. Further, these alterations were confirmed using independent ASCAT-based^13^ allele-specific copy number analysis (**Supplementary Figure 3**).

### Loss of heterozygosity events are preserved across primary, VC, and metastatic tissues

ASCAT-based copy number analysis enabled systematic identification of copy-neutral LOH events in addition to LOH associated with CN loss (**Supplementary Figure 3**). Across the cohort, LOH events of either type were most frequently observed on chromosomes 3, 9, and 14. Canonical chromosome 3 alterations occurred even more often as copy-neutral LOH (12 patients) than as heterozygous deletions (10 patients). Among patients with multiple sampled tissue sites, LOH profiles were generally consistent between anatomical compartments. In contrast, no LOH events were detected in benign tissues, underscoring the tumor-specific nature of these alterations.

### Cell culture-driven clonal remodeling in patient-derived cancer cells

To optimize PDC culture conditions and determine those that best preserve the genomic architecture of parental tumors, we first systematically evaluated multiple culture formats in a pilot cohort (P1, P4 and P7). For each case, PDCs were established under different conditions, including an irradiated mouse fibroblast feeder cell system (IR)^32^; 2D monolayers, both Matrigel-coated (MG) and Primaria adherent (PRIM); and 3D matrices of different concentration, including Matrigel (MG20) and GrowDex hydrogel (GD02 or GD04), tested in different supplemented media formulations (F-media vs BM).

Light microscopy images of PDCs cultured in Matrigel-coated dishes demonstrated adherent growth with morphology consistent with ccRCC, while preserving both interpatient and intrapatient heterogeneity (**Supplementary Figure 4A**). To confirm retention of characteristic molecular features, immunostaining was performed for established ccRCC-associated markers, including Ki-67 and CAIX (**Figure 3A**), as well as p-mTOR, β-catenin, and p53 (**Supplementary Figure 4B**). All examined PDC models expressed these markers, indicating the preservation of key signaling and proliferative characteristics. Ki-67 confirmed active proliferation across PDCs. CAIX, a canonical HIF target in ccRCC, was robustly expressed in all models, particularly in P4 and P1, suggesting maintenance of tumor-specific metabolic adaptations. The heterogeneous p-mTOR and β-catenin staining across samples reflected patient-specific differences. Nuclear p53 expression was observed in a subset of cells, with the strongest staining in P7, consistent with the presence of a highly clonal somatic *TP53* mutation in the corresponding tumor (**Supplementary Figure 4B** and **Figure 3B**). Collectively, these data demonstrate that PDC models preserve essential molecular features of ccRCC and provide biologically representative systems for downstream functional analyses.

**Figure 3.**
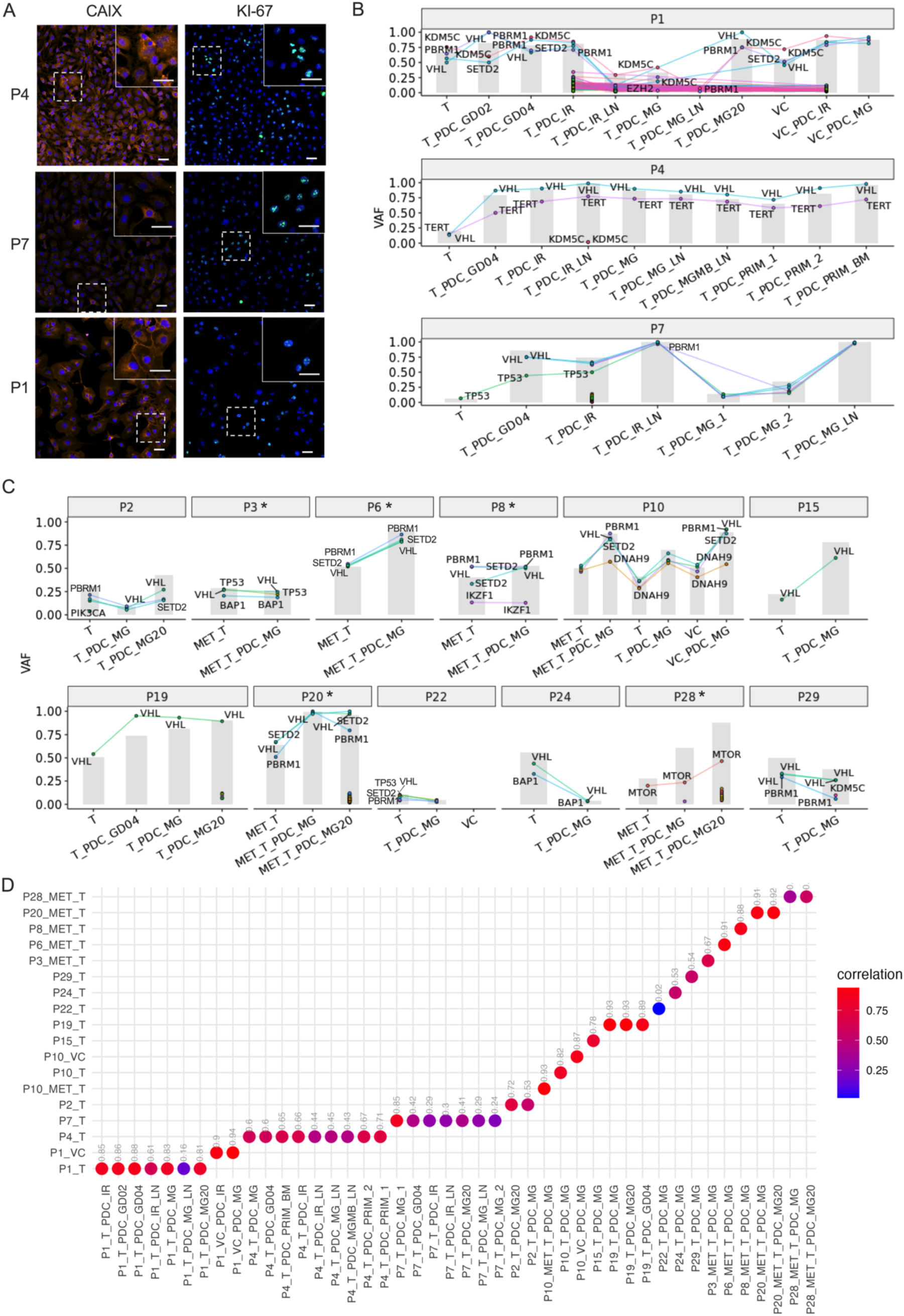
PDC models were confirmed to be representative by immunostaining and exome sequencing. **A.** Immunostaining of ccRCC markers, including Ki-67 and CAIX, in PDCs of P4, P7 and P1. Scale bars represent 50µm. **B-C.** VAFs of somatic mutations across tissues and corresponding representative PDC cultures. Each data point represents a distinct mutation, with connecting lines indicating within-patient VAF changes and concordance between tissues and corresponding PDCs across multiple specimens, including primary tumors, VC, and metastatic samples. All variants classified as clinically or functionally relevant by PCGR (Actionability Tier 1-2) are shown, although not every mutation is annotated with a gene symbol to maintain clarity. In addition, VHL, PBRM1, SETD2, DNAH9, KDM5C, and BAP1, are displayed with all coding mutations regardless of tier. Gray shading marks tumor cell content (“purity”) estimates for each sample. Asterisks (*) indicate patient samples from previously published datasets, reanalyzed and included here for comparative purposes. **D.** Genome-wide CNR Pearson correlations between primary tumor, VC, and metastatic tissue samples and their matched PDCs.

To determine the extent to which culture conditions preserve the genomic architecture of parental tumors, we performed WES-based validation of matched tissue-PDC pairs. Specifically, successful validation was defined as the retention of variants considered clinically actionable (Tier1-2 as annotated by PCGR^16^) or affecting established ccRCC-associated drivers *VHL, PBRM1, SETD2, DNAH9, KDM5C,* and *BAP1.* As shown in **Figure 3B**, 2D Matrigel-based cultures most faithfully retained key somatic driver mutations at VAFs comparable to matched tumor tissue. These findings indicate minimal clonal drift and effective tumor cell enrichment under these conditions. In contrast, 3D culture systems frequently yielded lower cell numbers and exhibited variable VAF shifts, consistent with selective clonal outgrowth or dilution effects, thereby limiting their suitability for high-throughput functional screening. In P1, culture on irradiated feeder cells resulted in a marked reduction in mutation VAFs, suggesting overgrowth of non-tumor components or selective clonal suppression. Additionally, feeder-based systems introduced non-human genomic background and increased contamination risk, occasionally leading to discordant VAF and CNA profiles that complicated downstream interpretation (e.g. note high number of likely artefactual low-VAF hits in IR samples of all three patients; **Figure 3B**). Collectively, these observations demonstrate that culture conditions can significantly influence clonal representation *in vitro*. Based on genomic fidelity, scalability, and throughput considerations, F-media on Matrigel-coated 2D plates was selected as the preferred platform for establishing low-passage ccRCC models.

### PDCs preserve ccRCC key markers and tumor-specific mutations

Among the 20 patients for whom PDC models were generated, at least one model was successfully validated by WES for 15 patients (75%). Models from these patients comprised primary tumors (n=10), VC tumor thrombi (n=3), and metastatic lesions (n=7), including five pancreatic metastases from our previously published DEDUCER subcohort study^7^, demonstrating reliable PDC establishment across clinically and anatomically distinct tissue sources (**Figure 3B** for pilot with diverse culture conditions and **Figure 3C** for subsequent patients). Together, these findings demonstrate that the majority of established PDC models maintain clinically and biologically relevant somatic driver alterations, supporting their suitability for downstream functional interrogation of genotype-phenotype relationships. For non-representative PDC models, their parental tissues typically displayed low driver mutation VAFs (0-0.3) and, by definition, lacked shared mutations with derived PDCs (**Supplementary Figure 5A**), consistent with insufficient tumor cell representation at baseline. To evaluate the broader genomic fidelity, we extended the analysis of the most frequent exonic mutations identified in parental tissues (**Figure 2A**) to include matched PDCs (**Supplementary Figure 6**). In most cases, the same mutations were detected in both tissue and PDC samples. However, subtle differences were observed in some models, reflecting shifts in clonal architecture during culture.

We next examined whether baseline tumor characteristics influenced successful establishment of genomically representative PDCs. Parent tissue VAFs of key driver mutations were compared between samples yielding representative versus non-representative PDC models (**Figure 3B-C and Supplementary Figure 5A-B**). For primary tumors and VC samples, no significant differences in driver mutation VAFs were observed (tumor: Wilcoxon p = 0.21; VC: p = 0.22). In contrast, metastatic tissues that successfully generated representative PDCs exhibited significantly higher driver mutation VAFs compared to those that did not (Wilcoxon p = 0.0036). Although metastatic disease status itself was not associated with representativeness (FET p = 1), this finding suggests that metastatic clones capable of successful *in vivo* expansion may also possess increased fitness for *in vitro* propagation. No significant associations were detected between representativeness and clinical parameters including age, sex, surgical approach, tumor necrosis, or tumor size. Nuclear grade was marginally higher in non-representative cases (Wilcoxon p = 0.059), whereas IMDC and Leibovich risk classifications showed no significant differences (**Supplementary Figure 5C**).

Taken together, these findings demonstrate that, when established under optimized conditions, the majority of PDC models preserve the mutational landscape of their parental tumors, including both dominant driver events and subclonal alterations. At the same time, the observed variability across individual cases highlights the influence of baseline tumor purity, clonal composition, and selective pressures during *in vitro* propagation. Importantly, successful establishment of genomically representative PDCs was more strongly associated with higher driver mutation burden in metastatic tissues than with conventional clinical parameters, suggesting that intrinsic clonal fitness rather than clinical risk category governs *in vitro* model fidelity. These results provide a robust genomic foundation for subsequent functional drug profiling while underscoring the need to account for culture-driven clonal dynamics when interpreting genotype-phenotype relationships.

### PDCs retain parental CNA landscapes

To assess whether PDC models recapitulate large-scale chromosomal architecture of their parental tumors, genome-wide CNR from matched tissue-PDC pairs were compared. Overall, high concordance was observed across the cohort, with strong genome-wide similarity between tumor samples and corresponding PDCs derived from primary tumors (mean Pearson r = 0.59), VC tumor thrombi (mean r = 0.9), and metastatic lesions (mean r = 0.77) (**Figure 3D**). Two exceptions were noted. P7 and P22 exhibited reduced correlation between parental tumor tissue and matched PDCs (P7 mean r = 0.44; P22 r = 0.02), indicating diminished preservation of copy number profiles in these cases. Both tumors displayed relatively low somatic driver mutation VAFs in the original tissue sample (**Figure 3B-C**), as well as quite low tumor cell content (P7 = 6.7% and P22 = 10.2%). For P7, given the high number of PDC models established with very consistent and ccRCC-typical CNA patterns (**Supplementary Figure 7**), the low correlation coefficient with the original tumor tissue is likely due to reduced parental CNA signals in a low tumor purity WES sample rather than lack of PDC genomic fidelity. Upon visual inspection, CNA patterns are indeed stable between all P7 PDC and the parental tissue samples, albeit the signal strength is notably reduced in the primary tumor (P7_T) and one PDC model (P7_T_PDC_MG_1) with lowest tumor cell content estimates (**Supplementary Figure 7** and **Figure 3B**). For P22, however, CNA profiles appear markedly distinct between the primary tumor and derived PDC model (**Supplementary Figure 7**). As P22 CNA profiles showed similar signal strength, and both the primary tumor and PDC had comparably low tumor cell content, increased intratumoral heterogeneity may have compromised faithful clonal representation during culture establishment. Notably, somatic driver alterations, including *VHL* mutations, were retained in the PDCs, albeit at low variant allele frequencies.

Although overall concordance was high, some degree of amplitude variation and regional divergences were observed (**Supplementary Figure 7**), consistent with subclonal selection during *in vitro* propagation. For example, P4-derived PDCs demonstrated more pronounced chromosomal gains and losses compared with the parental tumor, consistent with *in vitro* expansion of tumor clones and markedly higher tumor cell content in the resulting PDC vs primary tumor samples (mean tumor cell content of 83% vs 20%) (**Figure 3B**). A similar amplified copy number pattern was observed in P10 and was consistently retained across primary tumor, metastatic, and VC-derived PDCs, indicating stable clonal architecture across anatomical sites. In contrast, P24 harbored several alterations present in the tumor tissue which were not fully maintained in the corresponding PDCs. Notably, P24 PDCs were estimated to have lower tumor cell content (3.8%) than the matched primary tumor sample (56%), indicating that tumor CNA signals could have been simply diluted among an increased background of non-tumor cells. Across the cohort, recurrent ccRCC-associated chromosomal events were robustly preserved, including chromosome 3p loss in the majority of samples, prominent chromosome 5 gains in P1, P4, P7, P10 and P19, and chromosome 14q loss in models derived from P4, P7, P10, and P19. ASCAT-based allele-specific copy number analysis further showed that also total CN and LOH patterns were preserved in PDCs, while benign tissue–derived models retained their non-aberrant CN profiles (**Supplementary Figure 8-9**). Overall, these findings demonstrate that PDC models largely retain the genome-wide copy number landscape of their parental tumors, while also revealing sample-specific variability that likely reflects differences in tumor purity, clonal composition, and selective pressure during *in vitro* expansion.

### Functional drug screening identifies shared and patient-specific sensitivities in PDCs

We performed standardized DSRT on low-passage patient-derived cells (PDCs; mean passage 2.6) from primary tumors (T), adjacent benign tissue (B), VC tumor thrombi (VC), and metastases (MET) using a library of up to 535 oncology-relevant approved and investigational compounds in 384-well formats (**Figure 4A; Supplementary Tables 2-3**). Primary kidney epithelial cells and established renal cancer cell lines served as controls. DSRT screens corresponding to the same WES sample showed significant pairwise correlations (Pearson correlation coefficient 0.37-0.90, p < 0.05) and were averaged for downstream analyses (**Supplementary Figure 10A**). Specifically, we observed substantial variation in average DSS across patients’ PDCs, a pattern that persisted after adjusting for tissue of origin (Kruskal-Wallis p < 0.001; **Figure 4B** and **Supplementary Figure 10B**), though residual variability in the averages may also reflect tested drug library size or 2D/3D assay context (e.g. larger number of more effective drugs among larger drug sets).

**Figure 4.**
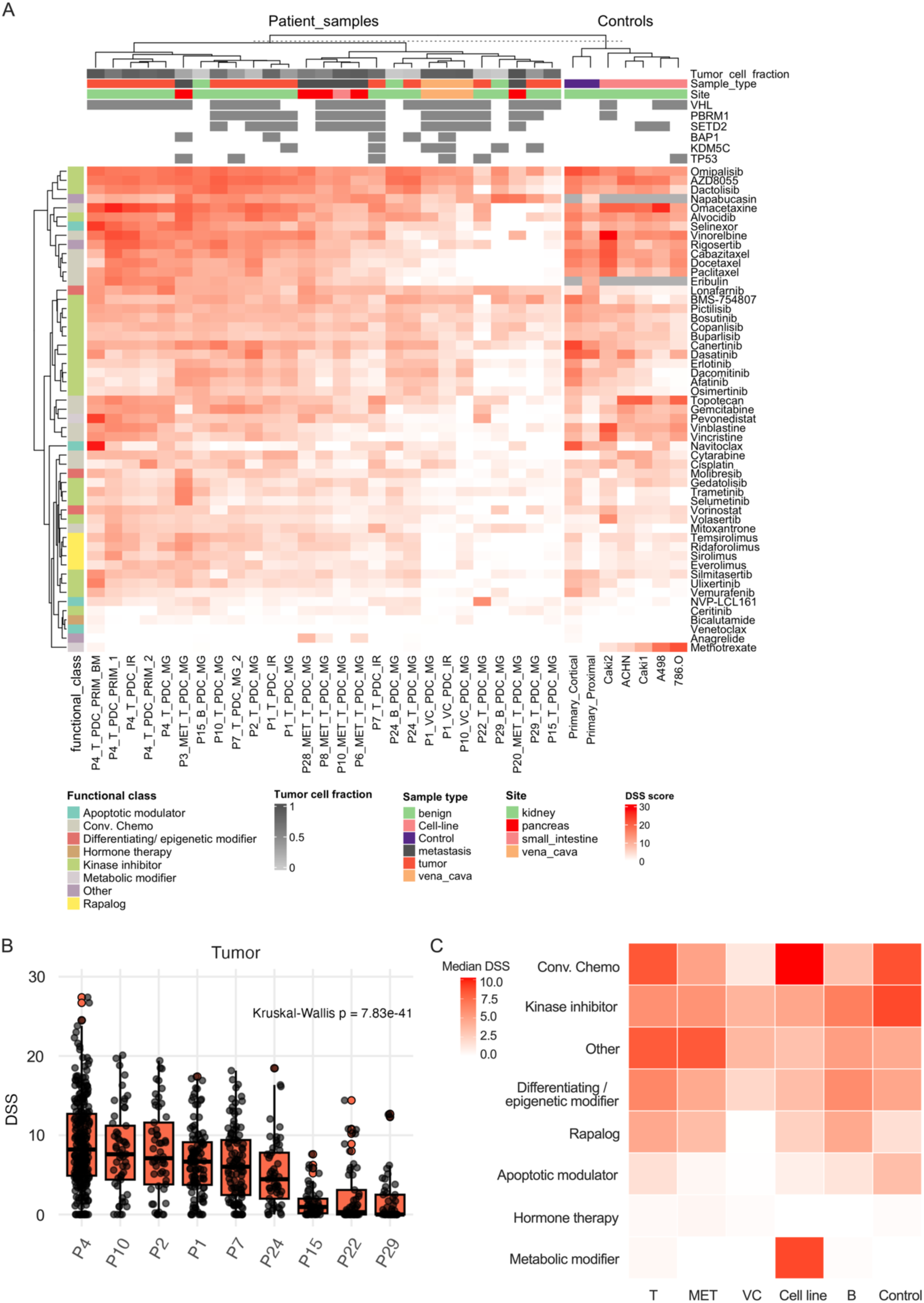
Functional drug sensitivity profiling of ccRCC PDC models. **A.** Drug sensitivity scores of representative PDC models for the drugs exhibiting mean DSS > 10 in at least one sample. Tumor cell fraction estimates, drug plate, passage, sample type, site and either presence or absence of commonly observed ccRCC mutations are indicated at the top of the heatmap while the functional drug class of each drug is indicated on the left. All DSRT replicates were averaged (mean) per matching WES sample (including replicates screened in 2D/3D conditions with FS2A (116 drugs)/FO5A (528 drugs) plates. For P1, P15 and P24, overall lower DSS values were observed in 3D- than 2D-DSRT). **B.** Per-patient drug sensitivity in tumor samples, ordered by descending median DSS. Each boxplot summarizes DSS values across compounds with mean DSS > 10 in at least one sample. **C.** Median DSS across T, MET, VC, cell line, B, and control (primary kidney epithelial cells) samples and functional drug classes.

The VC PDCs revealed a lower average DSS than PDCs derived from primary tumor/metastatic tissues (n=2 patients; Wilcoxon p<0.001) (**Supplementary Figure 10C**). Differential drug sensitivity analysis between primary tumor and VC samples identified 14 compounds with significantly increased sensitivity in primary tumor samples (delta DSS > 5, adjusted p-value<0.05), including omacetaxine, vinorelbine, alvocidib, cytarabine, and gemcitabine (**Supplementary Figure 10D**). These results suggest that drugs effective in tumor samples may exhibit reduced sensitivity in VC samples. Across all sample types, conventional chemotherapies and kinase inhibitors demonstrated the broadest efficacy, whereas hormone therapies showed minimal effectiveness (**Figure 4C**).

Cancer cell lines exhibited a distinctly stronger response to conventional chemotherapies than our PDCs, as well as unique sensitivity to metabolic modifier methotrexate. The top five most sensitive drugs for the tumor samples, on the other hand, were AZD8055, omacetaxine, omipalisib, vinorelbine, and dactolisib (**Supplementary Figure 10E**). Per-patient pairwise comparisons of matched 2D and 3D DSRT revealed that culture dimensionality influenced drug response in the majority of cases: five of eleven patients had significantly lower DSS in 3D cultures, two showed significantly lower DSS in 2D cultures, and four showed no significant difference between conditions (paired Wilcoxon signed-rank test, p < 0.05; **Supplementary Figure 11**). Collectively, DSRT in ccRCC PDCs delineated partially shared functional vulnerabilities alongside pronounced patient, site and culture-dependent drug sensitivities. Cohort-level analyses further revealed systematic differences between PDCs and cancer cell lines, underscoring the added value of PDC models for capturing clinically relevant therapeutic heterogeneity.

### Limited association between known genomic biomarkers and drug response

To determine whether established molecular biomarkers predict *ex vivo* drug sensitivity, we assessed associations between genomic alterations, including somatic SNVs, insertions and deletions (InDels), and CNAs, as annotated using PCGR^16^, and DSS derived from our DSRT. PCGR identified previously reported predictive genomic biomarkers associated with drug response in 27 of 28 patients (96%, all except P5) (**Supplementary Table 5**), suggesting actionability at the DNA level. In 9/27 patients, at least one of the identified biomarkers had been reported specifically as having clinical relevance in ccRCC (*VHL* and *BAP1 mutation*); the others had been reported in the context of other renal (*MET* amplification) or non-renal malignancies (all other biomarker alterations). However, integration of genomic annotations with functional drug response data revealed limited concordance. Among the drugs linked to these annotated biomarkers, effective *ex vivo* responses (DSS ≥ 5 in at least one sample) were observed for ten biomarker/drug pairs (**Supplementary Figure 12**). Importantly, drug sensitivity did not differ significantly between samples with and without the respective biomarker mutation or CNA, indicating that these alterations did not robustly predict functional response in our cohort.

### Novel genomic biomarkers are preserved in local but not distant disease

Given the limited concordance between the established genomic biomarkers and functional drug response, we next sought to identify novel molecular determinants of therapeutic sensitivity. We systematically tested associations between gene-level somatic mutations or copy number alterations and drug sensitivity scores across the cohort. This analysis yielded 19 significant DSS/CNA associations and no significant DSS/VAF associations (**Supplementary Table 6**), suggesting that copy number variation rather than mutation allele frequency was the dominant genomic correlate of *ex vivo* drug response in this dataset.

To determine whether these genotype-phenotype relationships were stable across anatomical contexts, we next separately evaluated the directionality of each significant drug/gene association in primary tumor (T), local VC tumor thrombi and distant metastatic (MET_T) samples. Associations, as quantified by linear regression coefficient (slope), were highly concordant between T and VC samples, with 12 of 13 drug/gene pairs demonstrating consistent direction of effect (e.g., higher copy number ratio associated with higher DSS in both T and VC samples). In contrast, associations were frequently reversed in MET_T samples, with only 1 of 7 drug/gene pairs maintaining concordant directionality (FET comparing T/VC vs T/MET_T agreement p = 0.001; **Figure 5A** and **Supplementary Figure 13-15**). Consistently, association slopes were strongly correlated between T and VC samples (Spearman rho = 0.71) but inversely correlated between T and MET_T samples (rho = -0.64) (**Figure 5B)**.

**Figure 5.**
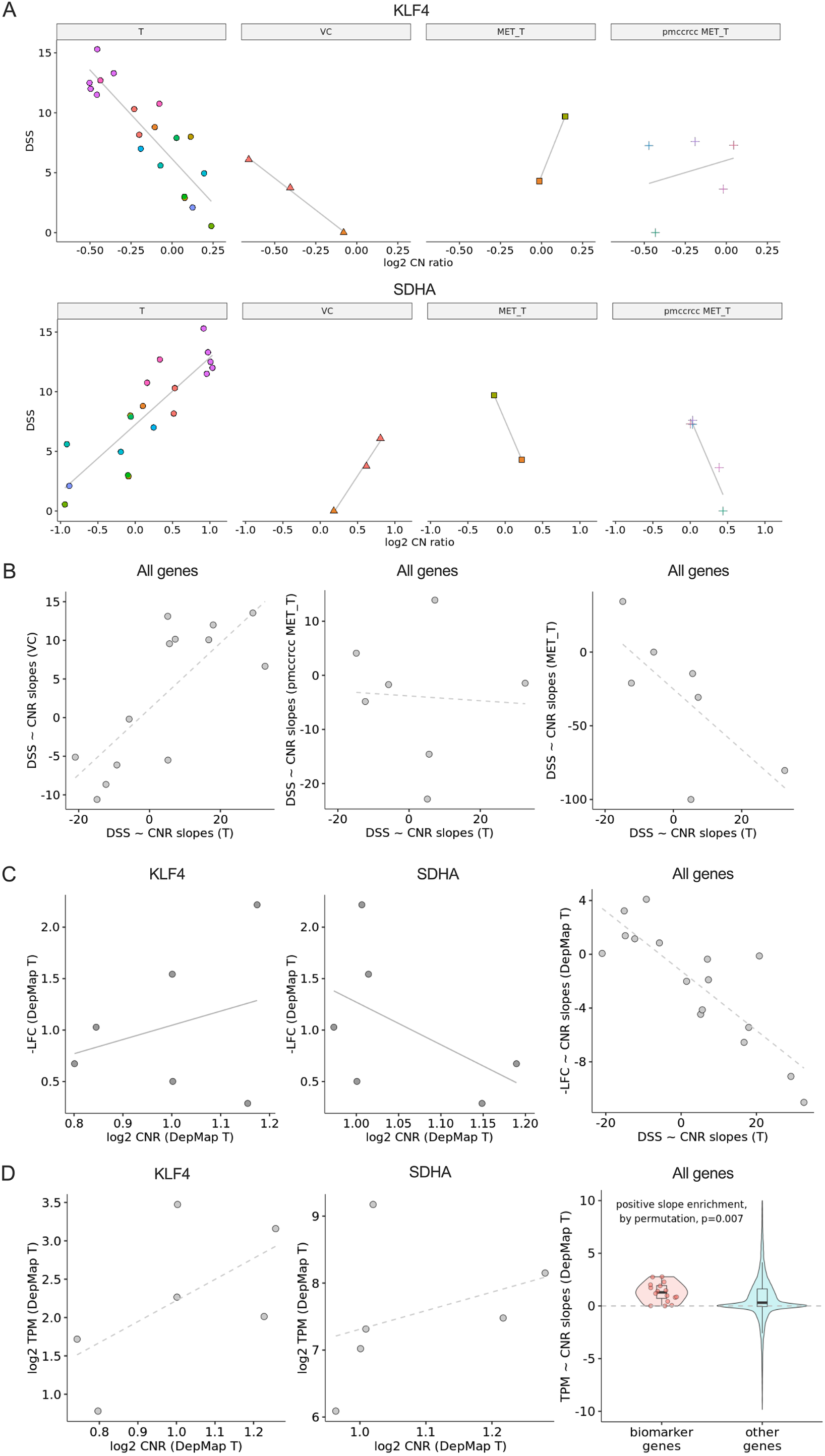
Gene-drug associations and copy number–linked sensitivity patterns across tumor contexts and DepMap validation datasets. **A.** Exemplary DSS ∼ CN ratio associations of Topotecan ∼ KLF4 (top) and Topotecan ∼ SDHA (bottom). Linear associations and trend lines are shown separately for primary tumor (T), vena cava (VC), non-pmccRCC metastatic (MET_T), and pmccRCC metastatic (pmccRCC MET_T) samples. While T and VC samples tend to follow the same trend, metastatic samples do not. **B.** Scatterplots comparing trend line slopes of DSS∼CN ratio associations between all genes for T/VC (left), T/pmccRCC MET_T (center), and T/MET_T (right). **C.** Exemplary -log2 fold change (-LFC) ∼ CNR ratio associations of Topotecan ∼ KLF4 (left), Topotecan ∼ SDHA (center), and all genes (right) using data from DepMap for six primary tumor derived cancer cell lines. **D.** Exemplary log2 TPM ∼ CNR ratio associations in the DepMap T dataset for KLF4 (left) and SDHA (center) genes, and slopes of all biomarker genes vs all other genes from the DepMap dataset (right). The latter shows that biomarker genes are significantly enriched for positive associations between genomic copy number and mRNA expression.

This metastasis-specific decoupling was further evaluated in an independent validation cohort comprising five pancreatic metastatic ccRCC (pmccRCC) samples not included in the discovery analysis^7^ (**Figure 5A** and **Supplementary Figures 13 and 15**). Only 3 of 7 drug/gene pairs with available DSRT data showed concordant directionality relative to primary tumor associations (FET comparing T/VC vs T/pmccRCC MET_T agreement p = 0.03). Moreover, regressions slopes were not correlated with those observed in primary tumors (Spearman rho = 0.11; Fisher’s z transformation test for independent correlations p = 0.09; **Figure 5B**), reinforcing the observation that genotype-drug response relationships are not preserved in distant metastatic contexts.

In addition, we examined the same significant drug/gene associations using *ex vivo* drug response and copy number data from six primary ccRCC-derived cancer cell lines in the DepMap PRISM Repurposing 24Q2 dataset (KMRC1, KMRC3, KMRC20, 786O, OSRC2, 769P)^23,24^. Despite an origin from primary ccRCC tumors, all 16 evaluable drug/gene pairs exhibited reversed directionality relative to our primary tumor associations (0/16 concordant), with strong inverse correlation of association slopes (Spearman rho = -0.81) (**Figure 5C** and **Supplementary Figure 13-14**). Lastly, we checked whether biomarker gene copy number was associated with, and potentially acting through, mRNA expression levels in the DepMap dataset^23,24^. Indeed, the 16 biomarker genes showed significant enrichment for genes with positive associations between DNA copy number and mRNA expression levels (15/16 genes, p=0.007, based on 1000 random gene set permutations; **Figure 5D** and **Supplementary Figure 16**).

To provide mechanistic context for the identified response-associated genes (**Supplementary Table 6** and **Supplementary Table 7**), we assessed their connectivity to corresponding drug targets within prior-knowledge signaling networks (**Supplementary Table 7**). All response-associated genes were connected to their respective targets through an average of 1.2 signaling intermediates (range 0-2). The Topotecan subnetwork exemplified this relationship, linking CDH17, ESR1, KLF4 and SDHA to TOP1 through short OmniPath signaling paths involving one or two intermediate genes (**Supplementary Figure 17**). While these connections do not imply causality, they contextualize the biomarkers’ associations with the particular drug response.

## Discussion

This study establishes a prospective functional precision medicine framework for ccRCC, integrating low-passage patient-derived cell models, drug testing, genomic profiling, and longitudinal clinical annotation. We demonstrate that while genomic annotation identifies potentially actionable alterations in the vast majority of patients, established DNA-based biomarkers explain only a limited proportion of *ex vivo* drug response variability. Moreover, we show that genotype-drug response associations identified in primary tumors and VC tumor thrombi are frequently disrupted in distant metastatic samples and immortalized ccRCC cell lines. Together, these findings reveal context-dependent decoupling of genotype and therapeutic phenotype and support the integration of functional drug testing with molecular profiling in ccRCC.

Our cohort recapitulated the canonical genomic landscape of ccRCC^31,33–37^, with VC thrombi and metastatic samples showing no significant differences from the primary tumor, supporting its relevance for downstream functional analyses. Phenotypic and genotypic characterization further demonstrated that optimized 2D Matrigel-based culture conditions enable the propagation of tumor-specific features in the majority of PDC models. Although minor shifts in variant allele frequencies and copy number amplitudes were observed, consistent with known culture-induced clonal bottlenecks, core driver alterations and chromosome-level LOH patterns were largely preserved. Importantly, successful establishment of representative PDCs was more strongly associated with higher tumor cell content and clonal dominance than with conventional clinical risk parameters, suggesting that intrinsic clonal fitness governs *in vitro* model fidelity.

Functional drug screening indicated partially shared vulnerabilities among patients, most notably to kinase inhibitors targeting the PI3K-AKT-mTOR axis. Our findings underscore that pathway-level dependencies in ccRCC are not uniform, and patient context can influence therapeutic phenotype^31,34^. Strikingly, although PCGR-based annotation identified putative actionable alterations in 96% of patients, *ex vivo* drug response could not be consistently predicted for any of the ten biomarker/drug pairs with decent activity in the DSRT. Thus, genomic actionability did not translate into functional predictability. This disconnect highlights a central limitation of single-locus, tier-based biomarker frameworks in tumor types where canonical, directly targetable drivers are rare, a principle increasingly demonstrated across solid tumors and hematologic malignancies^38^. Consistent with this, reduced drug sensitivity observed in VC samples was not explained by genomic features alone and may instead reflect higher variant allele frequencies or differences in transcriptomic states, as reported previously^39,40^.

Using linear mixed-effects modeling with leave-one-patient-out cross validation, we identified robust copy number-based genotype-drug associations. These associations were largely preserved between primary tumors and local VC tumor thrombi but were frequently reversed in distant metastatic samples and in primary tumor-derived immortalized cell lines from DepMap. The inverse correlation of association slopes suggests systematic genotype-phenotype rewiring during metastatic evolution and long-term *in vitro* adaptation. These findings imply that biomarker rules derived from metastatic tissue or established cell lines may not faithfully reflect primary tumor biology in ccRCC.

To provide mechanistic context for the drug response-associated genes, we examined their connectivity to the corresponding drug targets within prior-knowledge signaling networks. The resulting subnetworks recapitulated biological processes consistent with the known mechanisms of action of the respective compounds, including DNA damage/TOP1 inhibition (SN-38 and Topotecan), proteostasis stress (Ixazomib and Luminespib), apoptosis signaling (NVP-LCL161), and oncogenic or metabolic pathways^41–46^. In addition, certain high-priority markers are known to be involved in the pathways that are central to ccRCC biology, including mitochondrial metabolism (*S*DHA) and RTK/MAPK signaling (PTPN11)^47,48^. Taken together, the genomic association and network analyses provide complementary support for the biological interpretability of the identified drug response associations. Network analysis linked drug response-associated biomarkers to signaling pathways connected to established drug targets, providing mechanistic context without implying causality. However, these findings remain hypothesis-generating and require experimental and independent cohort validation to establish biological relevance and biomarker potential, particularly for cytotoxic agents such as Topotecan and SN-38, as proliferative state may influence drug sensitivity in PDC models and limit its ability to reflect responses in advanced ccRCC tumors.

From a translational perspective, our results support moving beyond genomics only toward integrative models that include multi-tissue sampling, high-fidelity PDCs, and functional drug response data. Structured molecular tumor board reporting systems remain essential for standardization and regulatory compliance, but their outputs should be interpreted within the broader biological and functional context of each tumor. Particularly in ccRCC, where pathway-level dysregulation predominates and directly actionable mutations are uncommon, functional testing may provide critical orthogonal information to refine therapeutic prioritization, as well as drug repurposing and discovery opportunities.

Several limitations warrant consideration. While PDC models are increasingly used in translational oncology, ensuring their translational reliability will require systematic evaluation and validation of PDC-derived drug responses against patient outcomes in clinical trials, including testing with clinically available therapies^49^. First, culture-induced clonal shifts may bias functional readouts away from the full breadth of intratumoral heterogeneity, potentially underrepresenting minor subclones relevant for resistance. Second, 2D assays may over- or underestimate drug sensitivity compared to 3D or *in vivo* contexts^50–52^. Third, established cancer cell lines showed distinct response patterns relative to PDCs and primary cells for some agents (e.g. methotrexate) as well as decoupled genotype/drug response patterns, underscoring the risk of overgeneralizing from immortalized models. Fourth, biomarkers derived from copy number gains and low- or non-coding tier variants remain hypothesis-generating without orthogonal validation, and they require careful contextualization with zygosity and co-occurring alterations to refine biological plausibility. Further limitations include the absence of native tumor microenvironment (TME), including stromal, vascular, and immune components, which precludes the assessment of spatially resolved tumor-immune-stromal interactions known to define aggressive phenotypes and poor prognosis in localized ccRCC^53^. ccRCC in particular is characterized by a highly immune-infiltrated microenvironment and is especially responsive to immune checkpoint blockade. Consequently, conventional *ex vivo* PDC models cannot capture immune-mediated therapeutic effects or dynamic tumor-immune interactions, including baseline immune states and treatment-induced microenvironment-driven resistance mechanisms within the RCC TME^54^. Instead, our system is well suited for interrogating tumor cell-intrinsic vulnerabilities but does not account for interaction-driven responses to immunotherapies or stromal signaling cues that may influence treatment outcomes *in vivo*.

In conclusion, we demonstrate that a functional precision oncology pipeline for ccRCC is feasible within a prospective clinical framework and provides clinically meaningful information beyond genomic annotation alone. Genotype-drug response relationships are context-dependent, preserved between primary tumors and local tumor thrombi but frequently disrupted in distant metastases and immortalized models. Integrating functional drug testing with genomic profiling offers a scalable strategy to account for interpatient and intersite heterogeneity and may improve therapeutic prioritization in ccRCC.

## Declarations

### Ethics approval and consent to participate

This study was approved by the ethics committee of the Hospital District of Helsinki and Uusimaa (HUS) (Dnro 154/13/03/02/2016, 15.03.2017; HUS/850/2017, 14.07.2021). All patients provided written informed consent prior to inclusion in the study. The study was conducted in accordance with the Declaration of Helsinki and all relevant regulations, including the General Data Protection Regulation (GDPR). Patient material and clinical data were collected and processed under the Institutional Ethical Review Board-approved protocol.

### Consent for publication

Written informed consent for publication was obtained from all participants included in this study. No identifying personal information is included in the manuscript or supplementary materials.

### Data availability

The whole-exome sequencing data supporting the conclusions of this article are not publicly available due to sensitivity of patient information but are available from the corresponding author upon reasonable request. These data are stored in controlled-access infrastructure at CSC - IT Center for Science. Clinical data and drug sensitivity data are provided in Supplementary Tables 1 and 2. Public validation datasets were obtained from DepMap (25Q3 and 24Q2) https://depmap.org/portal/data_page/?tab=allData and cBioPortal https://www.cbioportal.org/study/summary?id=kirc_tcga.

### Author contributions

MF, TJL, RK, and VP conceptualized the study. MF, PM, RM, and MA performed the laboratory investigations. TJL and RK conducted the formal bioinformatics analyses with support from LJG and SP and curated the data. PP, MM, MR-M, HS, PK, PJ, TM, and AR were responsible for clinical sample and data acquisition. VC, MG, OK, AR, and VP provided resources. MF, TJL, RK and VP wrote the original draft of the manuscript. MF, TJL, RK, MP, PM, MA, ER, MR-M, PK, MG, AR, and VP contributed to data interpretation, manuscript review, and editing. Visualization was performed by TJL, RK, and MF. The study was supervised by VP and AR Project administration was carried out by MF and VP, and funding was acquired by VC, AR and VP. All authors read and approved the final manuscript.

### Competing interests

O.K. is a cofounder and board member of Sartar Therapeutics (not directly relevant to this study). The authors declare no other conflicts of interest.

## Acknowledgements

We acknowledge FIMM High Throughput Biomedicine Unit (HTB) supported by Helsinki Institute of Life Science-HiLIFE, Biocenter Finland, and EU-OPENSCREEN, FIMM High Content Imaging and Analysis (HCA) Unit supported by HiLIFE, University of Helsinki, Biocenter Finland and EuroBioImaging, and FIMM HiPREP and Genomics core units supported by HiLIFE and Biocenter Finland. Library preparation, sequencing and DRAGEN analysis were performed by the FIMM Genomics NGS unit at University of Helsinki supported by HiLIFE and Biocenter Finland. We thank Single-Cell Technologies (SCT, Hungary) for providing the software used for microscopic image formatting. The authors wish to acknowledge CSC – IT Center for Science, Finland, for generous computational resources. We greatly acknowledge all the patients participating in the study.

## Funding

The following grants are reported during the study: the Finnish Cultural Foundation (MF), Orion Research Foundation sr (MF), Munuaissäätiö (MF), Kymenlaakso Foundation of Finnish Cultural Foundation (VP), Jane and Aatos Erkko Foundation (AR), the Research Council of Finland (AR, VP/grant 373151, iCAN Flagship), the Data and AI-Enhanced Molecular Medicine (DAIMM) Research Initiative, funded through a Brain Gain grant from the Sigrid Jusélius Foundation, the Jane and Aatos Erkko Foundation, and the Finnish Medical Foundation (ER, OK), Cancer Foundation Finland (AR, VP), Governmental research funding (AR), Astra Zeneca (AR), and FICAN South (AR, VP). The funders had no role in study design, data collection, or analysis.

## Additional material

**Supplementary Table 1**, File format:.xls, Title/description of the data: Clinical information of ccRCC patients included in the study cohort.

**Supplementary Table 2**, File format:.xls, Title/description of the data: Low-passage PDC cultures and culture conditions used in DSRT experiments.

**Supplementary Table 3**, File format:.xls, Title/description of the data: Approved and investigational compound library used for drug screening.

**Supplementary Table 4**, File format:.xls, Title/description of the data: Gene mutation and frequency data across TCGA and DEDUCER tumor, VC, and metastatic samples.

**Supplementary Table 5**, File format:.xls, Title/description of the data: Patient- and sample-level PCGR biomarker annotations including genomic alterations and clinical actionability in ccRCC.

**Supplementary Table 6**, File format:.xls, Title/description of the data: DSS associations with CNA and VAF in ccRCC samples.

**Supplementary Table 7,** File format:.xls, Title/description of the data: List of direct molecular targets for biomarker-associated drugs.

**Supplementary Figures 1-17**, File format:.pdf, Title/description of the data: Additional figures and figure legends, supporting the findings described in the manuscript.

## Notes

### Summary of Updates

This version of the manuscript has been revised to add per-patient pairwise comparison of drug sensitivity scores between matched 2D and 3D DSRT assays (See new Supplementary Figure 11) and to update the mechanistic context for the identified drug response-associated genes (see new Supplementary Fig. 17 and Supplementary Table 7). Accordingly, Materials and Methods, Results and Discussion sections have been updated. Authors and affiliations have been updated.

