## Supplementary Figures for "Site-Dependent Decoupling of Drug-Biomarker Associations in Clear Cell Renal Cell Carcinoma Revealed by Functional Profiling of Patient-Derived Cell Models"

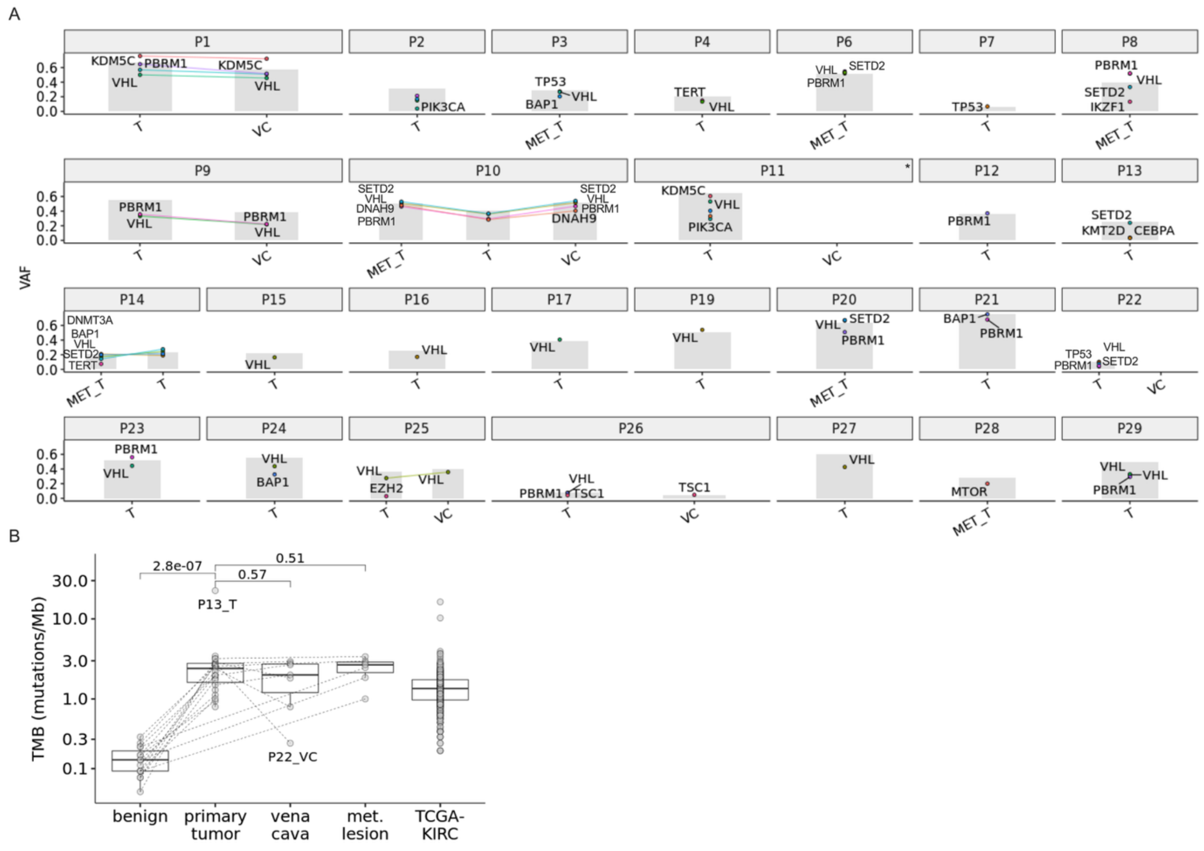


***Supplementary Figure 1. Genomic characterization of ccRCC and PM-ccRCC tissue samples.***

***A.*** *VAFs of selected somatic mutations across 27 patient samples. P5 is omitted, as no somatic mutations of clinical actionability (PCGR Tier1-2) or known canonical drivers VHL, PBRM1, SETD2, DNAH9, KDM5C and BAP1 were identified in this individual. Each data point represents a distinct mutation. Identical mutations found in multiple samples of a given patient are connected by a line to visualize changes in VAF. Not every mutation is labeled with a gene symbol to retain legibility. Gray shading indicates estimated tumor cell content of each sample. Asterisks (*) indicate patient samples from previously published datasets, reanalyzed and included here for comparative purposes.* ***B.*** *Boxplot of TMB in 23 primary ccRCC tumors, 7 VC samples, 7 metastatic samples, 15 benign samples and 360 primary ccRCC tumors from the TCGA-KIRC cohort for reference. TMB was calculated as the total number of non-silent somatic coding variants per sample. No significant differences were* *observed between primary tumors and VC (p = 0.57) or metastatic samples (p = 0.51) using an unpaired Wilcoxon test, while benign samples did have significantly lower TMB (p = 2.8e^-7).*

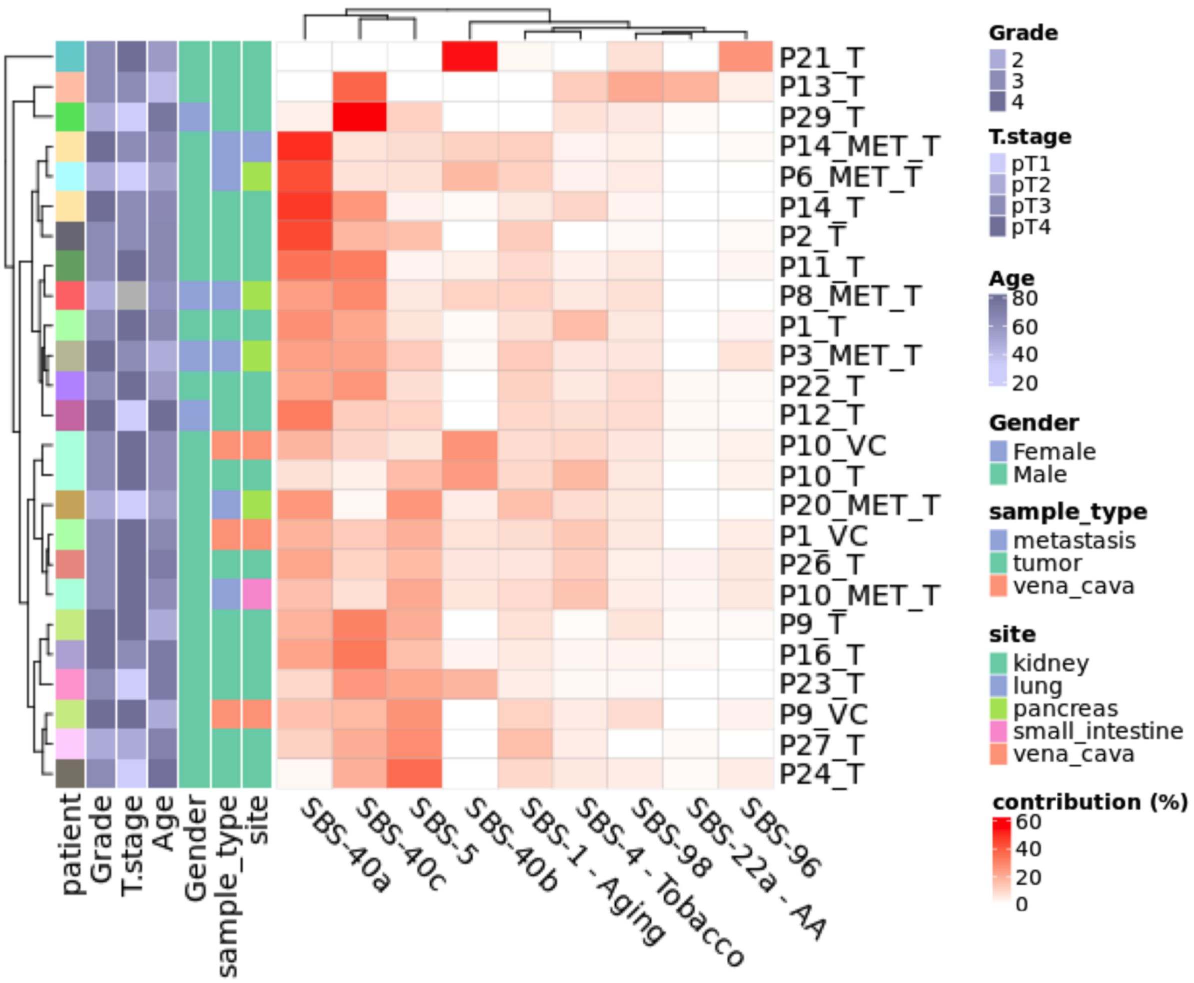


***Supplementary Figure 2. Mutational Signature Architecture of ccRCC Tumors.***

*Heatmap of somatic single base substitution (SBS) mutational signatures (n = 20/28). Color intensity denotes the percent contribution of each SBS signature to the total SBS mutation burden in each sample (e.g. aristolochic acid (AA) exposure related SBS-22a contributes ~20% of total SBSs in P13_T). Samples with <200 SBS events were excluded from this analysis because signatures could not be deconvoluted reliably.*

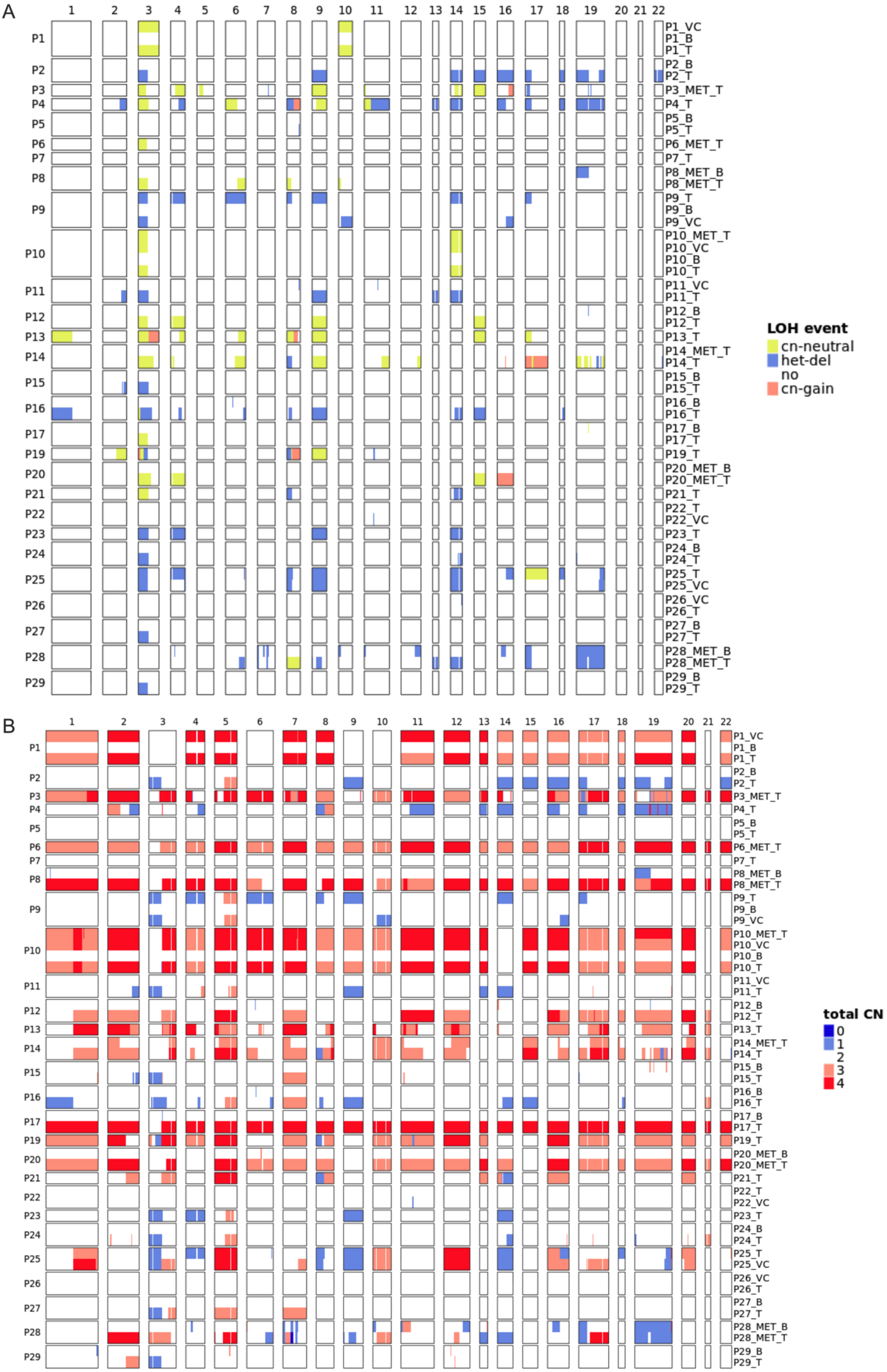
A
B
LOH event
cn-neutral
het-del
no
cn-gain
total CN
0
1
2
3
4

***Supplementary Figure 3. Cohort-wide ASCAT analysis of copy number and LOH alterations in ccRCC.***

***A.*** *ASCAT-derived CN and loss-of-heterozygosity (LOH) events across 28 patients. Recurrent LOH events, particularly affecting chromosome 3, were observed across the cohort, with variable patterns of CN–neutral LOH and heterozygous deletions, highlighting interpatient and intrapatient genomic heterogeneity.* ***B.*** *Genome-wide total CN profiles determined by ASCAT are shown for benign, tumor, VC, and metastatic samples across all patients.*

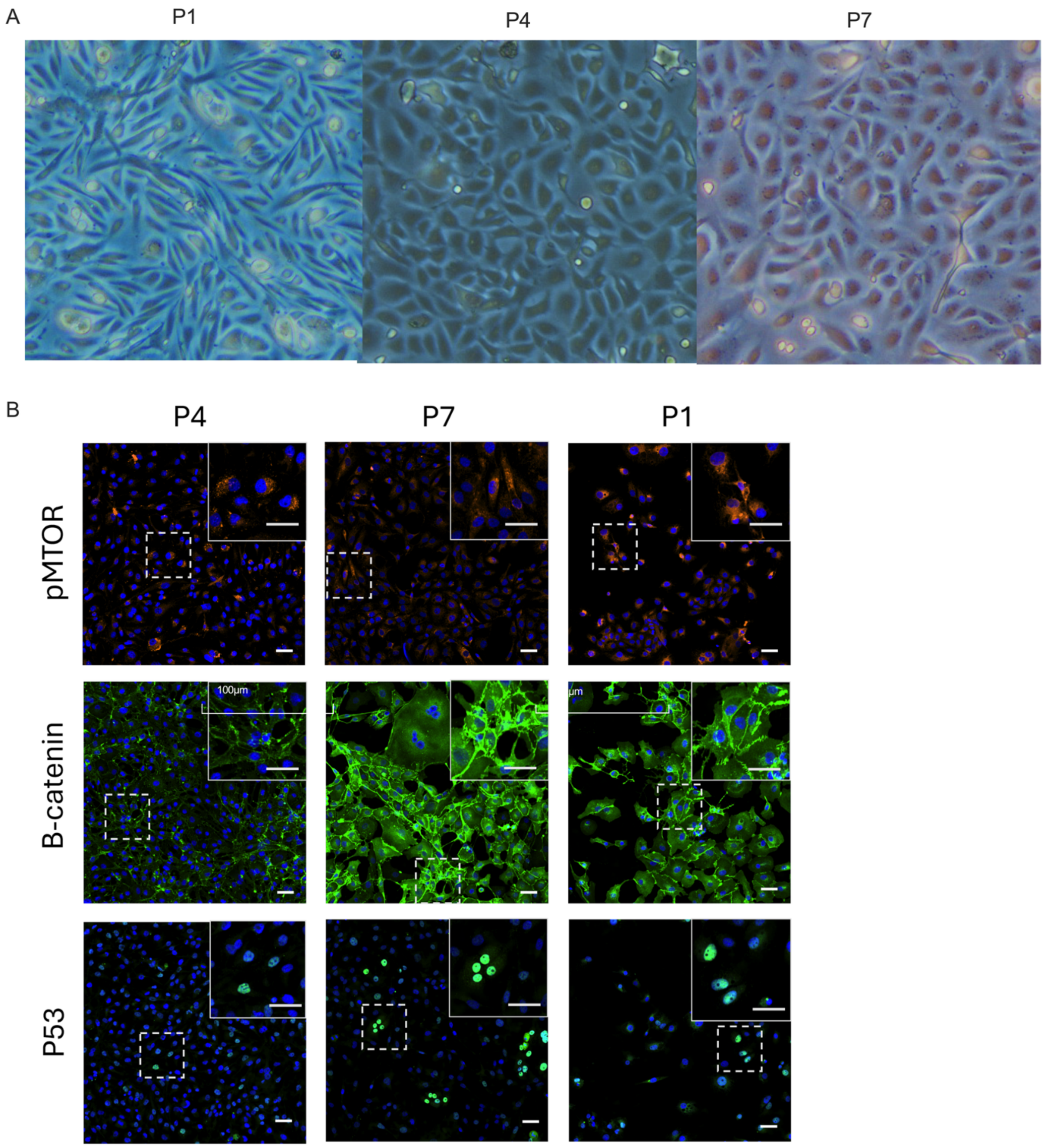


***Supplementary Figure 4. Morphology and marker validation of early passage ccRCC PDCs.*** ***A.*** *PDCs from P1, P4 and P7 cultured on matrigel-coated dishes at passage 1. Scale bars represent 100 µm.* ***B.*** *IF staining of PDCs from three ccRCC patients (P4, P7, and P1) with zoomed-in regions highlighting common ccRCC markers, including* p-mTOR*,* β-catenin*, and* p53*. Scale bars represent 50 µm.*

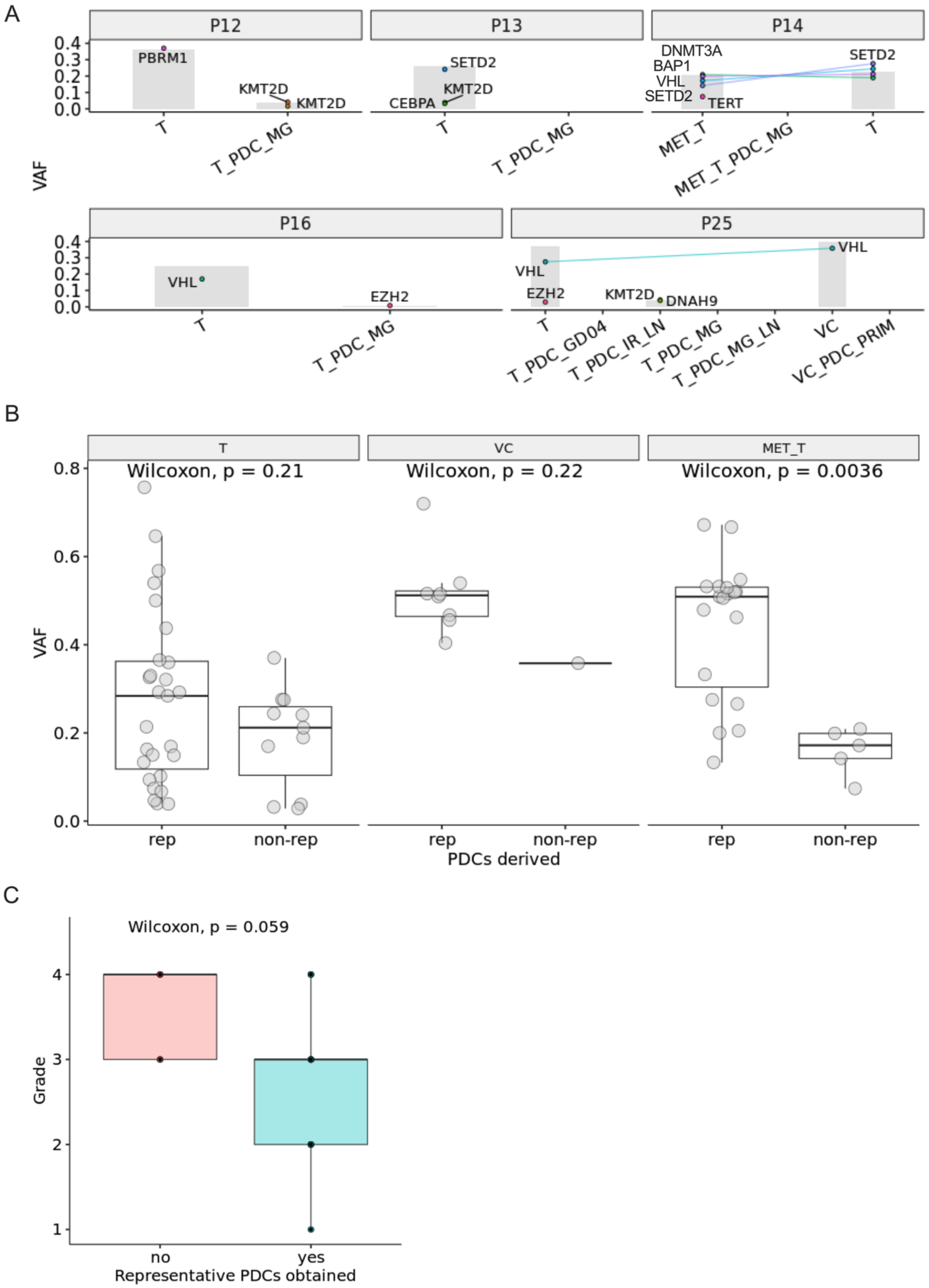
A
P12
P13
P14
VAF
0.4
0.3
0.2
0.1
0.0
PBRM1
KMT2D
KMT2D
SETD2
KMT2D
CEBPA
DNMT3A
BAP1
VHL
SETD2
TERT
SETD2
T
T_PDC_MG
T
T_PDC_MG
MET_T
MET_T_PDC_MG
T
P16
P25
VHL
EZH2
VHL
EZH2
KMT2D
DNAH9
VHL
T
T_PDC_MG
T
T_PDC_GD04
T_PDC_IR_LN
T_PDC_MG
T_PDC_MG_LN
VC
VC_PDC_PRIM
B
T
VC
MET_T
Wilcoxon, p = 0.21
Wilcoxon, p = 0.22
Wilcoxon, p = 0.0036
VAF
0.8
0.6
0.4
0.2
0.0
rep
non-rep
rep
non-rep
rep
non-rep
PDCs derived
C
Wilcoxon, p = 0.059
Grade
4
3
2
1
no
yes
Representative PDCs obtained

***Supplementary Figure 5. Association of parental tumor genomics and grade with representative versus non-representative PDC models.***

***A.*** *VAF plots showing clinically relevant (PCGR Tier1-2) and recurrent ccRCC driver mutations in patients without representative PDCs. Each data point represents a distinct mutation, with connecting lines indicating variants shared across different samples of the same patient. All detected mutations occurred at low frequencies, generally below 30% VAF. None of the mutations detected in the parent tissues of the five patients (P12, P13, P14, P16, and P25) were present in the corresponding PDCs.* ***B.*** *Comparison of these clinically relevant/recurrent driver mutation VAFs in the parent tissues from which representative PDCs were derived (rep) or not derived (non-rep) shows significantly higher parental VAFs only in representative vs non-representative metastatic samples (Wilcoxon p = 0.0036).* ***C.*** *Correlation between tumor grade and representative versus non-representative PDC models.*

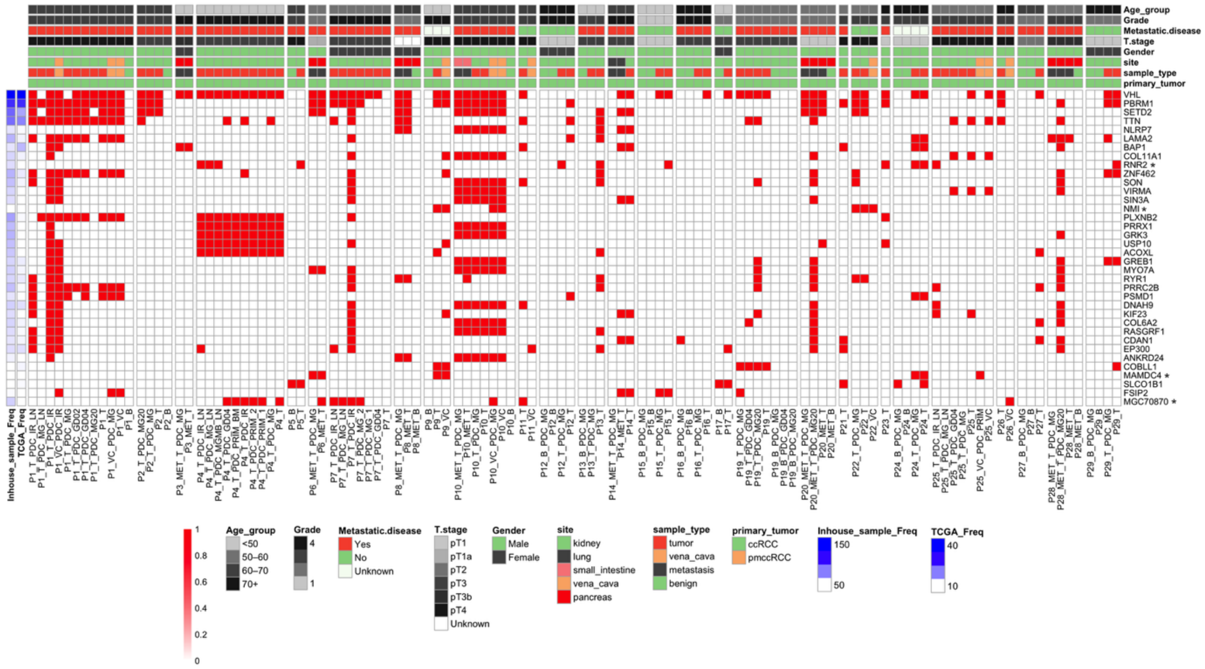

***Supplementary Figure 6. Preservation of mutational landscapes in ccRCC PDCs.***

*Retained mutations in ccRCC PDC models validated by WES. Mutations present in >3 tumor samples (same as Fig2A) shown in tumors and PDCs derived from them. PDCs of P20, P28, P3, P8, and P6 have been revisualized from previously published data sets for comparative purposes. Genes labeled with * were not found to be mutated in the TCGA dataset.*

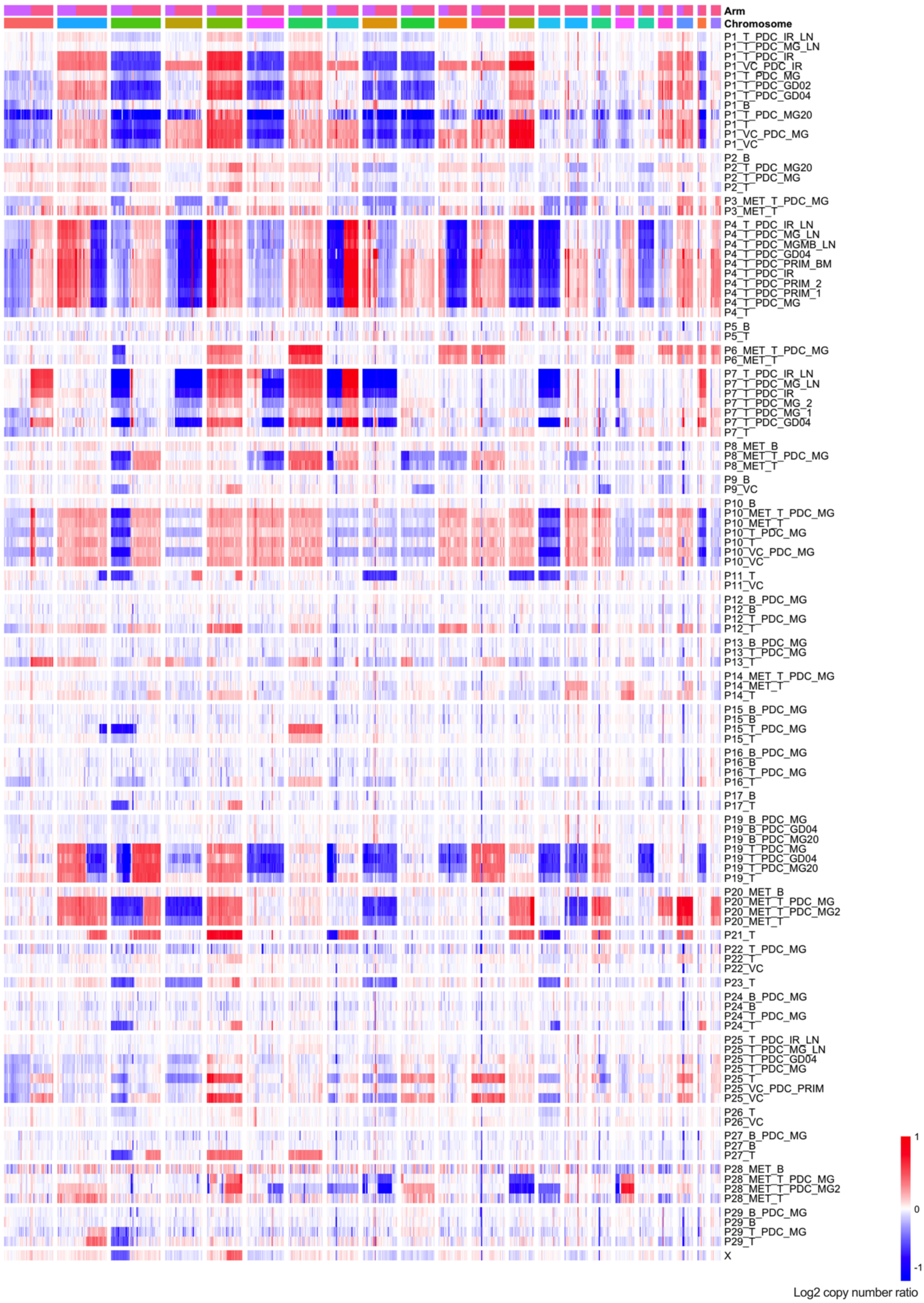

Arm
Chromosome
P1_T_PDC_IR_LN
P1_T_PDC_MG_LN
P1_T_PDC_IR
P1_VC_PDC_IR
P1_T_PDC_MG
P1_T_PDC_GD02
P1_T_PDC_GD04
P1_B
P1_T_PDC_MG20
P1_T
P1_VC_PDC_MG
P1_VC
P2_B
P2_T_PDC_MG20
P2_T_PDC_MG
P2_T
P3_MET_T_PDC_MG
P3_MET_T
P4_T_PDC_IR_LN
P4_T_PDC_MG_LN
P4_T_PDC_MGMB_LN
P4_T_PDC_GD04
P4_T_PDC_PRIM_BM
P4_T_PDC_IR
P4_T_PDC_PRIM_2
P4_T_PDC_PRIM_1
P4_T_PDC_MG
P4_T
P5_B
P5_T
P6_MET_T_PDC_MG
P6_MET_T
P7_T_PDC_IR_LN
P7_T_PDC_MG_LN
P7_T_PDC_IR
P7_T_PDC_MG_2
P7_T_PDC_MG_1
P7_T_PDC_GD04
P7_T
P8_MET_B
P8_MET_T_PDC_MG
P8_MET_T
P9_B
P9_VC
P10_B
P10_MET_T_PDC_MG
P10_MET_T
P10_T_PDC_MG
P10_T
P10_VC_PDC_MG
P10_VC
P11_T
P11_VC
P12_B_PDC_MG
P12_B
P12_T_PDC_MG
P12_T
P13_B_PDC_MG
P13_T_PDC_MG
P13_T
P14_MET_T_PDC_MG
P14_MET_T
P14_T
P15_B_PDC_MG
P15_B
P15_T_PDC_MG
P15_T
P16_B_PDC_MG
P16_B
P16_T_PDC_MG
P16_T
P17_B
P17_T
P19_B_PDC_MG
P19_B_PDC_GD04
P19_B_PDC_MG20
P19_T_PDC_MG
P19_T_PDC_GD04
P19_T_PDC_MG20
P19_T
P20_MET_B
P20_MET_T_PDC_MG
P20_MET_T_PDC_MG2
P20_MET_T
P21_T
P22_T_PDC_MG
P22_T
P22_VC
P23_T
P24_B_PDC_MG
P24_B
P24_T_PDC_MG
P24_T
P25_T_PDC_IR_LN
P25_T_PDC_MG_LN
P25_T_PDC_GD04
P25_T_PDC_MG
P25_T
P25_VC_PDC_PRIM
P25_VC
P26_T
P26_VC
P27_B_PDC_MG
P27_B
P27_T
P28_MET_B
P28_MET_T_PDC_MG
P28_MET_T_PDC_MG2
P28_MET_T
P29_B_PDC_MG
P29_B
P29_T_PDC_MG
P29_T
X
1
0
-1
Log2 copy number ratio

***Supplementary Figure 7. Chromosomal copy number profiles of PDCs and matched patient tissues.***

*Chromosomal-level CN analysis of PDC cultures and their matched tissue samples. CN profiles are shown for corresponding primary tumor, VC, and/or metastatic tissue where available, alongside the respective PDCs derived from the same patient. The analysis illustrates the degree of concordance in large-scale chromosomal gains and losses, highlighting the preservation of patient-specific CNAs in PDCs.*

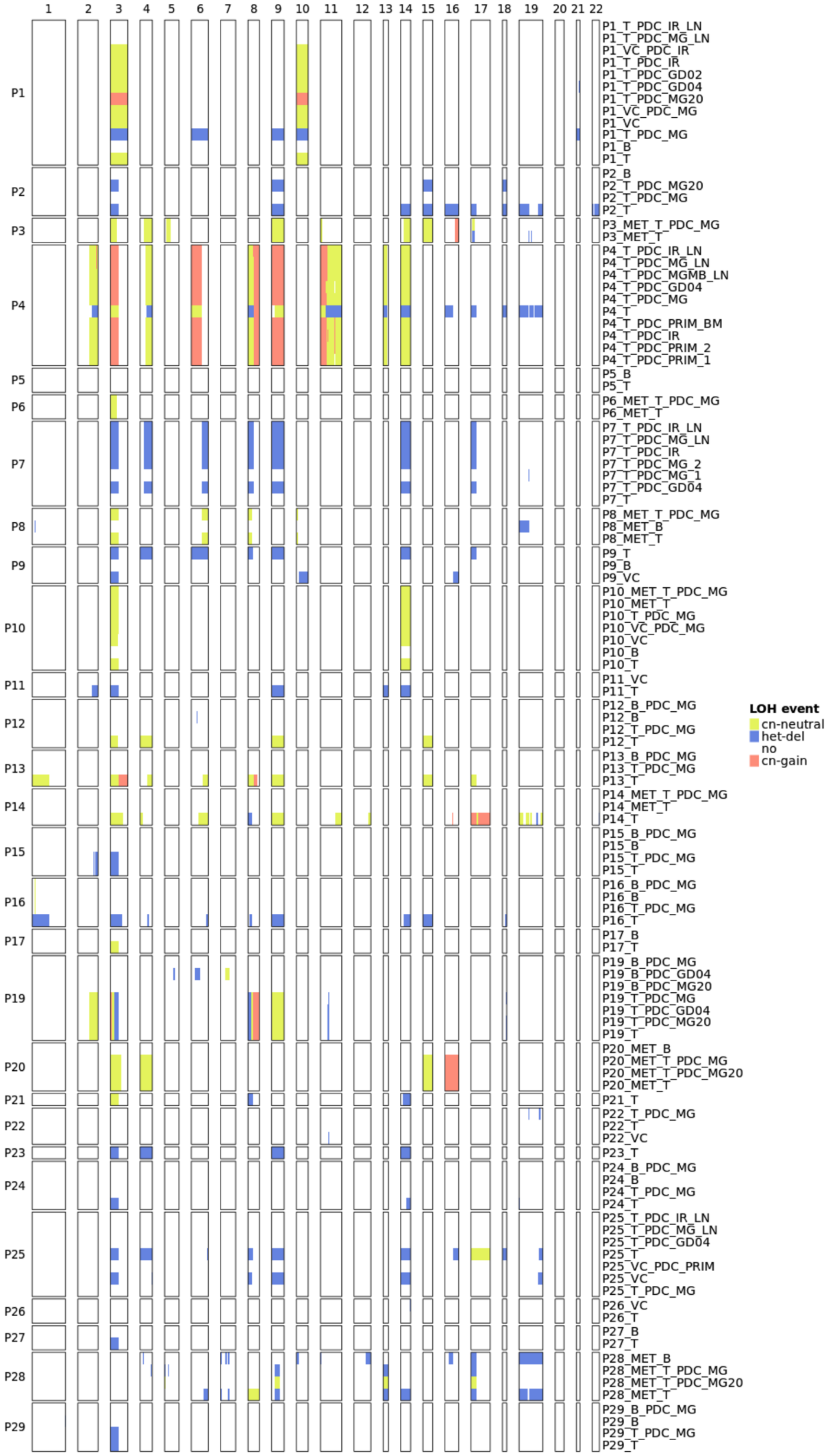
1 2 3 4 5 6 7 8 9 10 11 12 13 14 15 16 17 18 19 20 21 22
P1
P2
P3
P4
P5
P6
P7
P8
P9
P10
P11
P12
P13
P14
P15
P16
P17
P19
P20
P21
P22
P23
P24
P25
P26
P27
P28
P29
P1_T_PDC_IR_LN
P1_T_PDC_MG_LN
P1_VC_PDC_IR
P1_T_PDC_IR
P1_T_PDC_GD02
P1_T_PDC_GD04
P1_T_PDC_MG20
P1_VC_PDC_MG
P1_VC
P1_T_PDC_MG
P1_B
P1_T
P2_B
P2_T_PDC_MG20
P2_T_PDC_MG
P2_T
P3_MET_T_PDC_MG
P3_MET_T
P4_T_PDC_IR_LN
P4_T_PDC_MG_LN
P4_T_PDC_MGMB_LN
P4_T_PDC_GD04
P4_T_PDC_MG
P4_T
P4_T_PDC_PRIM_BM
P4_T_PDC_IR
P4_T_PDC_PRIM_2
P4_T_PDC_PRIM_1
P5_B
P5_T
P6_MET_T_PDC_MG
P6_MET_T
P7_T_PDC_IR_LN
P7_T_PDC_MG_LN
P7_T_PDC_IR
P7_T_PDC_MG_2
P7_T_PDC_MG_1
P7_T_PDC_GD04
P7_T
P8_MET_T_PDC_MG
P8_MET_B
P8_MET_T
P9_T
P9_B
P9_VC
P10_MET_T_PDC_MG
P10_MET_T
P10_T_PDC_MG
P10_VC_PDC_MG
P10_VC
P10_B
P10_T
P11_VC
P11_T
P12_B_PDC_MG
P12_B
P12_T_PDC_MG
P12_T
P13_B_PDC_MG
P13_T_PDC_MG
P13_T
P14_MET_T_PDC_MG
P14_MET_T
P14_T
P15_B_PDC_MG
P15_B
P15_T_PDC_MG
P15_T
P16_B_PDC_MG
P16_B
P16_T_PDC_MG
P16_T
P17_B
P17_T
P19_B_PDC_MG
P19_B_PDC_GD04
P19_B_PDC_MG20
P19_T_PDC_MG
P19_T_PDC_GD04
P19_T_PDC_MG20
P19_T
P20_MET_B
P20_MET_T_PDC_MG
P20_MET_T_PDC_MG20
P20_MET_T
P21_T
P22_T_PDC_MG
P22_T
P22_VC
P23_T
P24_B_PDC_MG
P24_B
P24_T_PDC_MG
P24_T
P25_T_PDC_IR_LN
P25_T_PDC_MG_LN
P25_T_PDC_GD04
P25_T
P25_VC_PDC_PRIM
P25_VC
P25_T_PDC_MG
P26_VC
P26_T
P27_B
P27_T
P28_MET_B
P28_MET_T_PDC_MG
P28_MET_T_PDC_MG20
P28_MET_T
P29_B_PDC_MG
P29_B
P29_T_PDC_MG
P29_T
LOH event
cn-neutral
het-del
no
cn-gain

***Supplementary Figure 8. Preservation of chromosome-level copy number and LOH patterns in representative ccRCC PDCs.***

*ASCAT-derived CN and LOH profiles across representative PDCs (**A**). Colors indicate CN status: yellow, CN–neutral LOH; light blue, heterozygous CN-loss; white, no CNA; and light red, CN gain of one or more copies. Representative PDCs largely preserve patient-specific chromosome-level LOH patterns observed in the corresponding parental tissues, including recurrent alterations affecting chromosome 3.*

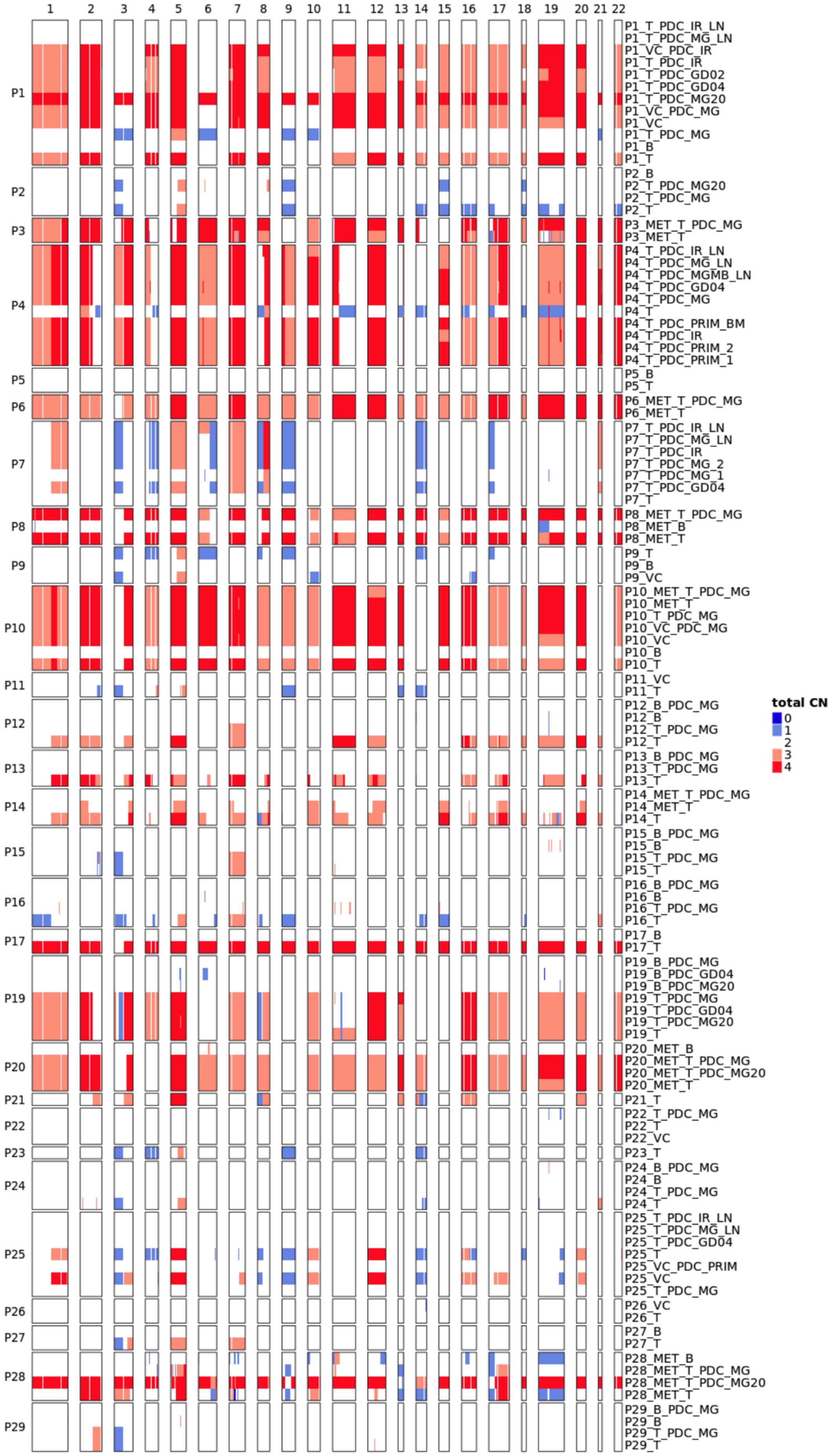

1 2 3 4 5 6 7 8 9 10 11 12 13 14 15 16 17 18 19 20 21 22
P1
P1_T_PDC_IR_LN
P1_T_PDC_MG_LN
P1_VC_PDC_IR
P1_T_PDC_IR
P1_T_PDC_GD02
P1_T_PDC_GD04
P1_T_PDC_MG20
P1_VC_PDC_MG
P1_VC
P1_T_PDC_MG
P1_B
P1_T
P2
P2_B
P2_T_PDC_MG20
P2_T_PDC_MG
P2_T
P3
P3_MET_T_PDC_MG
P3_MET_T
P4
P4_T_PDC_IR_LN
P4_T_PDC_MG_LN
P4_T_PDC_MGMB_LN
P4_T_PDC_GD04
P4_T_PDC_MG
P4_T
P4_T_PDC_PRIM_BM
P4_T_PDC_IR
P4_T_PDC_PRIM_2
P4_T_PDC_PRIM_1
P5
P5_B
P5_T
P6
P6_MET_T_PDC_MG
P6_MET_T
P7
P7_T_PDC_IR_LN
P7_T_PDC_MG_LN
P7_T_PDC_IR
P7_T_PDC_MG_2
P7_T_PDC_MG_1
P7_T_PDC_GD04
P7_T
P8
P8_MET_T_PDC_MG
P8_MET_B
P8_MET_T
P9
P9_T
P9_B
P9_VC
P10
P10_MET_T_PDC_MG
P10_MET_T
P10_T_PDC_MG
P10_VC_PDC_MG
P10_VC
P10_B
P10_T
P11
P11_VC
P11_T
P12
P12_B_PDC_MG
P12_B
P12_T_PDC_MG
P12_T
P13
P13_B_PDC_MG
P13_T_PDC_MG
P13_T
P14
P14_MET_T_PDC_MG
P14_MET_T
P14_T
P15
P15_B_PDC_MG
P15_B
P15_T_PDC_MG
P15_T
P16
P16_B_PDC_MG
P16_B
P16_T_PDC_MG
P16_T
P17
P17_B
P17_T
P19
P19_B_PDC_MG
P19_B_PDC_GD04
P19_B_PDC_MG20
P19_T_PDC_MG
P19_T_PDC_GD04
P19_T_PDC_MG20
P19_T
P20
P20_MET_B
P20_MET_T_PDC_MG
P20_MET_T_PDC_MG20
P20_MET_T
P21
P21_T
P22
P22_T_PDC_MG
P22_T
P22_VC
P23
P23_T
P24
P24_B_PDC_MG
P24_B
P24_T_PDC_MG
P24_T
P25
P25_T_PDC_IR_LN
P25_T_PDC_MG_LN
P25_T_PDC_GD04
P25_T
P25_VC_PDC_PRIM
P25_VC
P25_T_PDC_MG
P26
P26_VC
P26_T
P27
P27_B
P27_T
P28
P28_MET_B
P28_MET_T_PDC_MG
P28_MET_T_PDC_MG20
P28_MET_T
P29
P29_B_PDC_MG
P29_B
P29_T_PDC_MG
P29_T
total CN
0
1
2
3
4

***Supplementary Figure 9. Genome-wide ASCAT copy number profiles of representative ccRCC patient-derived cultures.***

*Genome-wide total ASCAT CN profiles of representative patient-derived ccRCC cultures. Multiple PDC models derived from tumor, VC, and metastatic tissues are shown. CN states are displayed on a scale from 0 to 4: dark blue = CN 0 (2 copies lost), light blue = CN 1 (1 copy lost), white = CN 2 (neutral), light red = CN 3 (one copy gained), and dark red = CN 4 (2 copies gained). Representative PDCs largely retain patient-specific CN gains and losses observed in the parental tissues, while a subset of models shows deviations consistent with subclonal selection during in vitro propagation.*

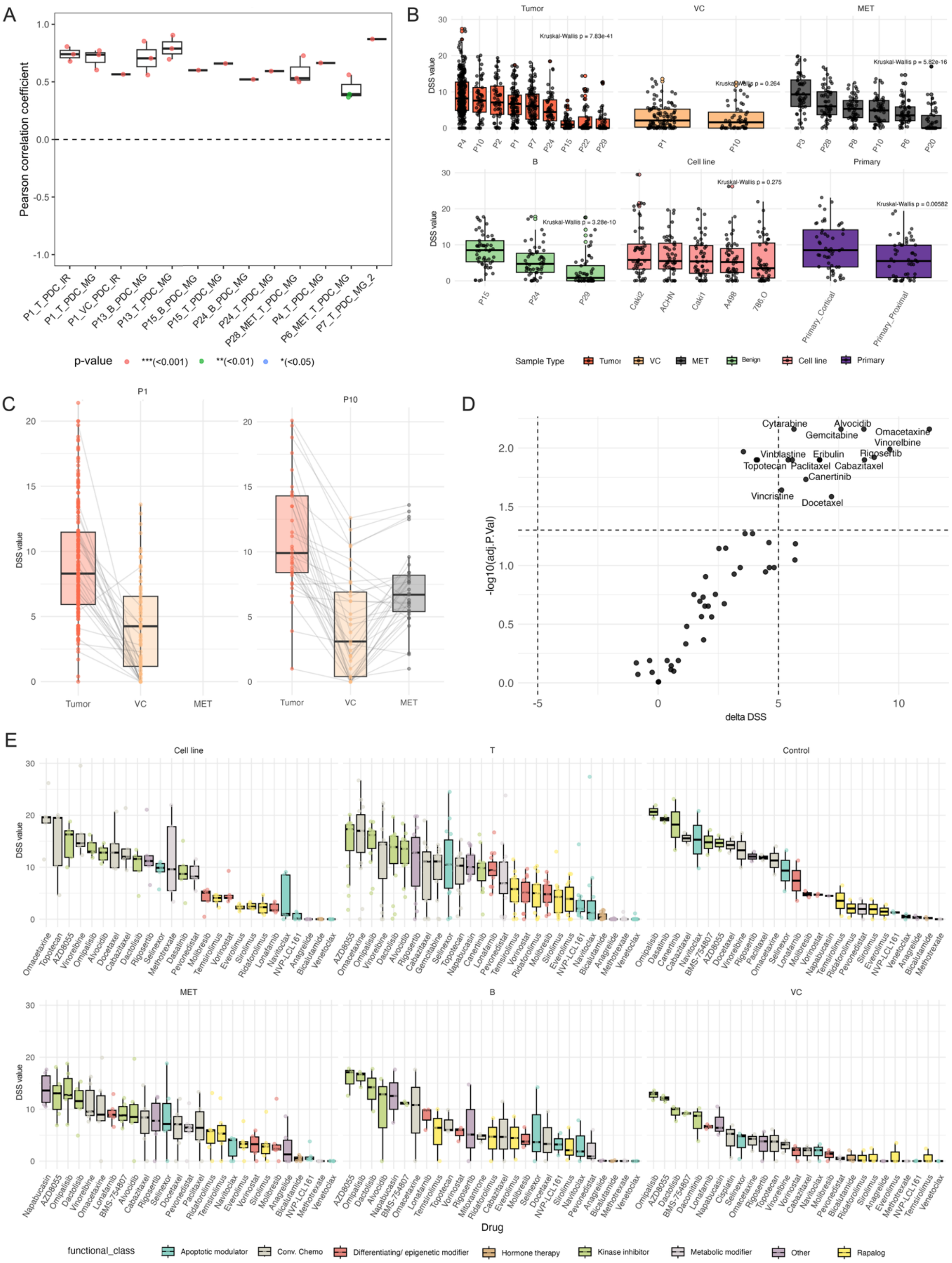
A
B
C
D
E
Pearson correlation coefficient
p-value
***(<0.001)
**(<0.01)
*(<0.05)
Tumor
VC
MET
B
Cell line
Primary
Kruskal-Wallis p = 7.83e-41
Kruskal-Wallis p = 0.264
Kruskal-Wallis p = 5.82e-16
Kruskal-Wallis p = 3.28e-10
Kruskal-Wallis p = 0.275
Kruskal-Wallis p = 0.00582
DSS value
Sample Type
Benign
P1
P10
-log10(adj.P.Val)
delta DSS
Cytarabine
Alvocidib
Gemcitabine
Omacetaxine
Vinorelbine
Vinblastine
Eribulin
Rigosertib
Topotecan
Paclitaxel
Cabazitaxel
Canertinib
Vincristine
Docetaxel
Control
T
Drug
functional_class
Apoptotic modulator
Conv. Chemo
Differentiating/ epigenetic modifier
Hormone therapy
Kinase inhibitor
Metabolic modifier
Other
Rapalog

***Supplementary Figure 10. Drug sensitivity profiles across sample types, patients, and assay conditions.***

***A.*** *Correlation of DSRT samples originating from the same WES sample.* ***B.*** *DSS per patient by sample type, sorted by descending median DSS. All DSRT replicates were included, including replicates screened in 2D/3D conditions and with FS2A (116 drugs)/FO5A (528 drugs) plates. For P1, P15 and P24, overall lower DSS values were observed in 3D- than 2D-DSRT).* ***C.*** *Boxplot comparing DSS across P1 and P10 PDC T, VC, and MET sample types.* ***D.*** *Differential drug sensitivity analysis of tumor vs. VC samples (delta DSS > 5, adjusted p < 0.05).* ***E.*** *Top 5 drugs per functional class in each sample group.*

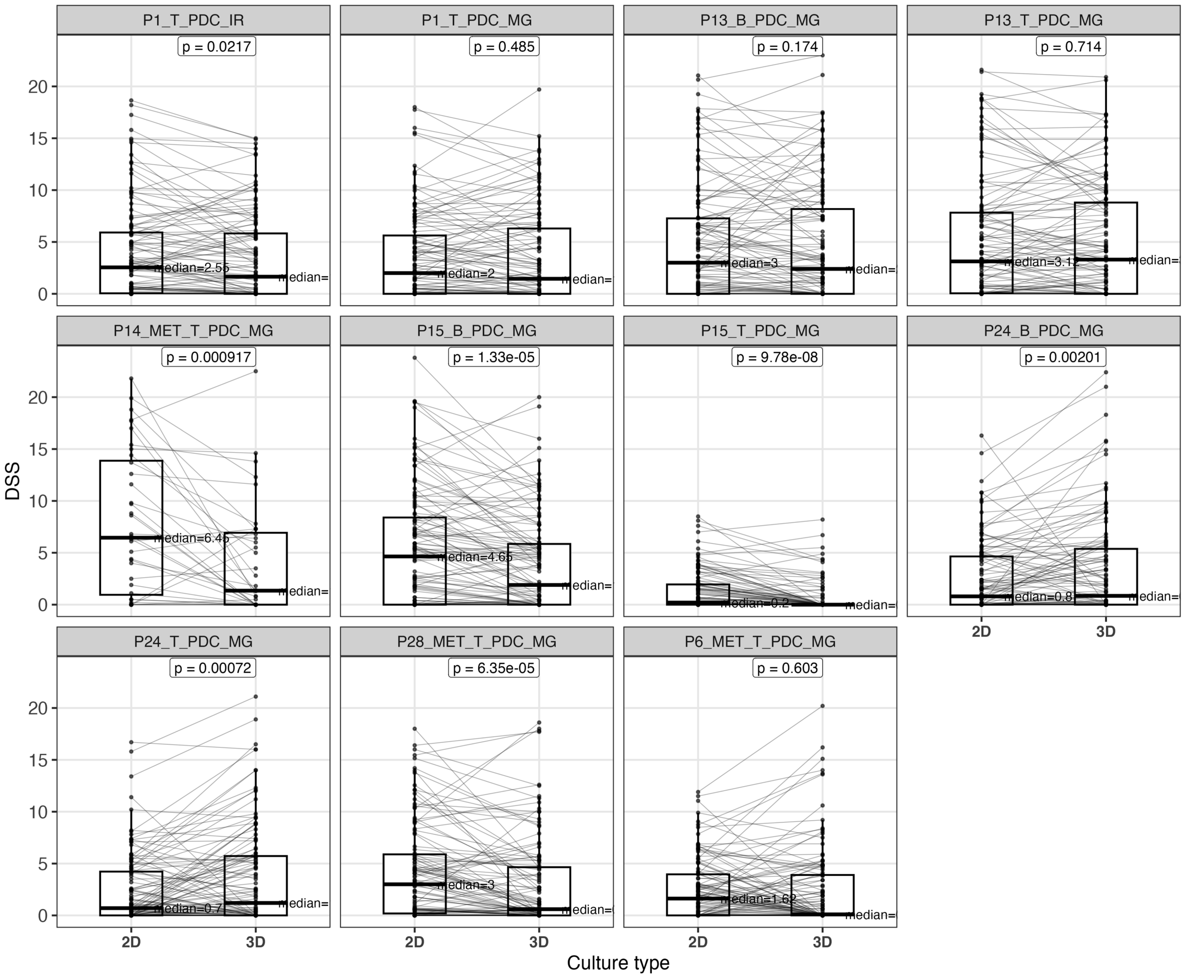


***Supplementary Figure 11. Per-patient pairwise comparison of drug sensitivity scores between matched 2D and 3D DSRT assays.***

*Pairwise comparison of DSS between matched 2D and 3D DSRT for each individual patient sample (n = 11). For each patient, DSS values obtained under 2D and 3D culture conditions in DSRT were compared using a paired Wilcoxon signed-rank test ($p < 0.05$). Five patient samples exhibited significantly lower DSS in 3D cultures (corresponding to higher DSS in 2D cultures), two patient samples showed significantly lower DSS in 2D cultures (corresponding to higher DSS in 3D cultures) and four patient samples showed no statistically significant difference between 2D and 3D cultures.*

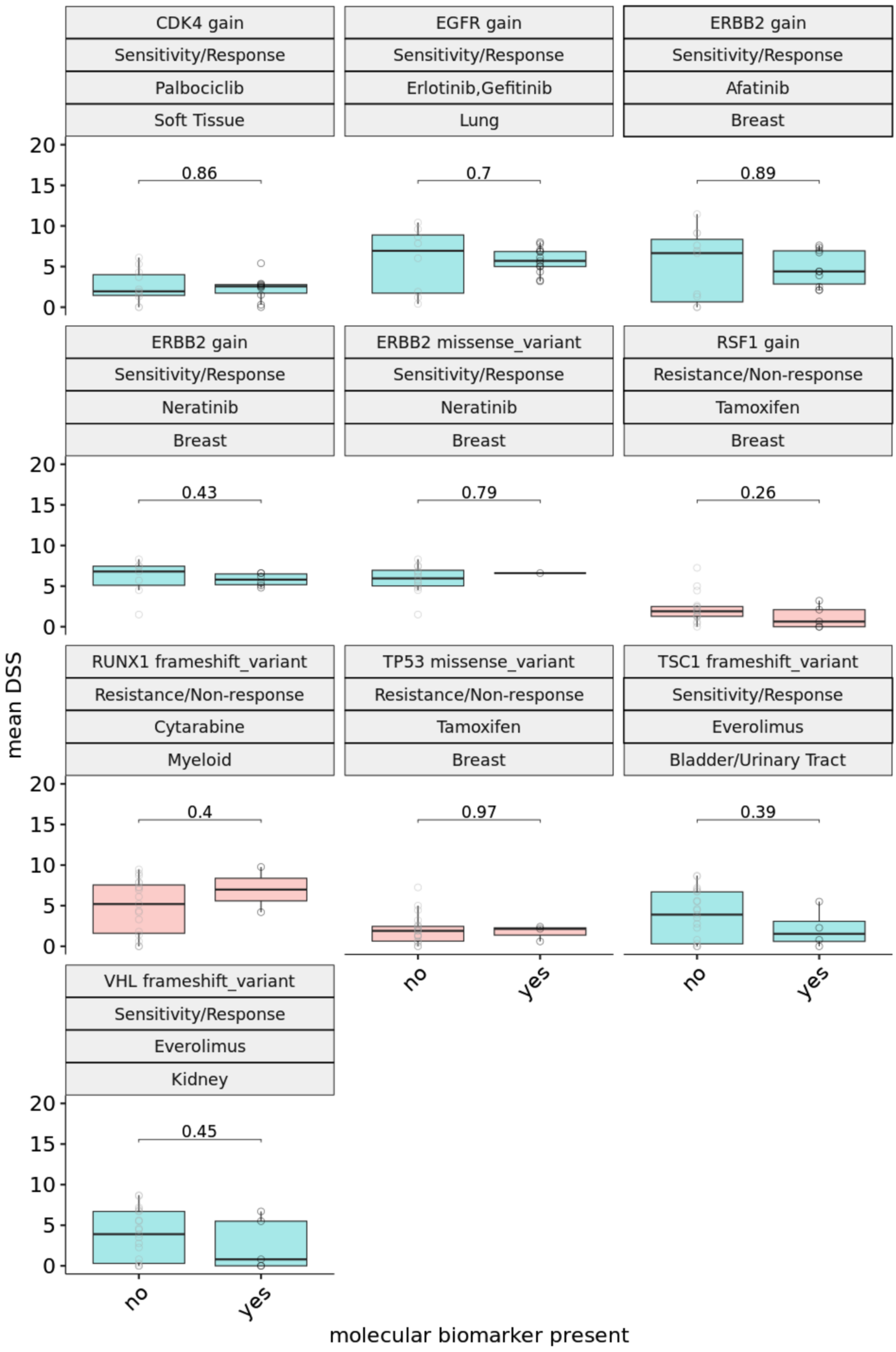

CDK4 gain
Sensitivity/Response
Palbociclib
Soft Tissue
0.86
EGFR gain
Sensitivity/Response
Erlotinib,Gefitinib
Lung
0.7
ERBB2 gain
Sensitivity/Response
Afatinib
Breast
0.89
ERBB2 gain
Sensitivity/Response
Neratinib
Breast
0.43
ERBB2 missense_variant
Sensitivity/Response
Neratinib
Breast
0.79
RSF1 gain
Resistance/Non-response
Tamoxifen
Breast
0.26
RUNX1 frameshift_variant
Resistance/Non-response
Cytarabine
Myeloid
0.4
TP53 missense_variant
Resistance/Non-response
Tamoxifen
Breast
0.97
TSC1 frameshift_variant
Sensitivity/Response
Everolimus
Bladder/Urinary Tract
0.39
VHL frameshift_variant
Sensitivity/Response
Everolimus
Kidney
0.45
0
5
10
15
20
no
yes
mean DSS
molecular biomarker present

***Supplementary Figure 12. Known genomic biomarker associations with ex vivo drug response in ccRCC PDCs.***

*Ten previously established genomic biomarkers for which alterations were detected in our ccRCC PDCs and associated drugs tested in our DSRT, with DSS >=5 in at least one sample. None of them show a significant difference in DSS between samples with and without the respective biomarker alteration (Wilcoxon p ranging from 0.26-0.97).*

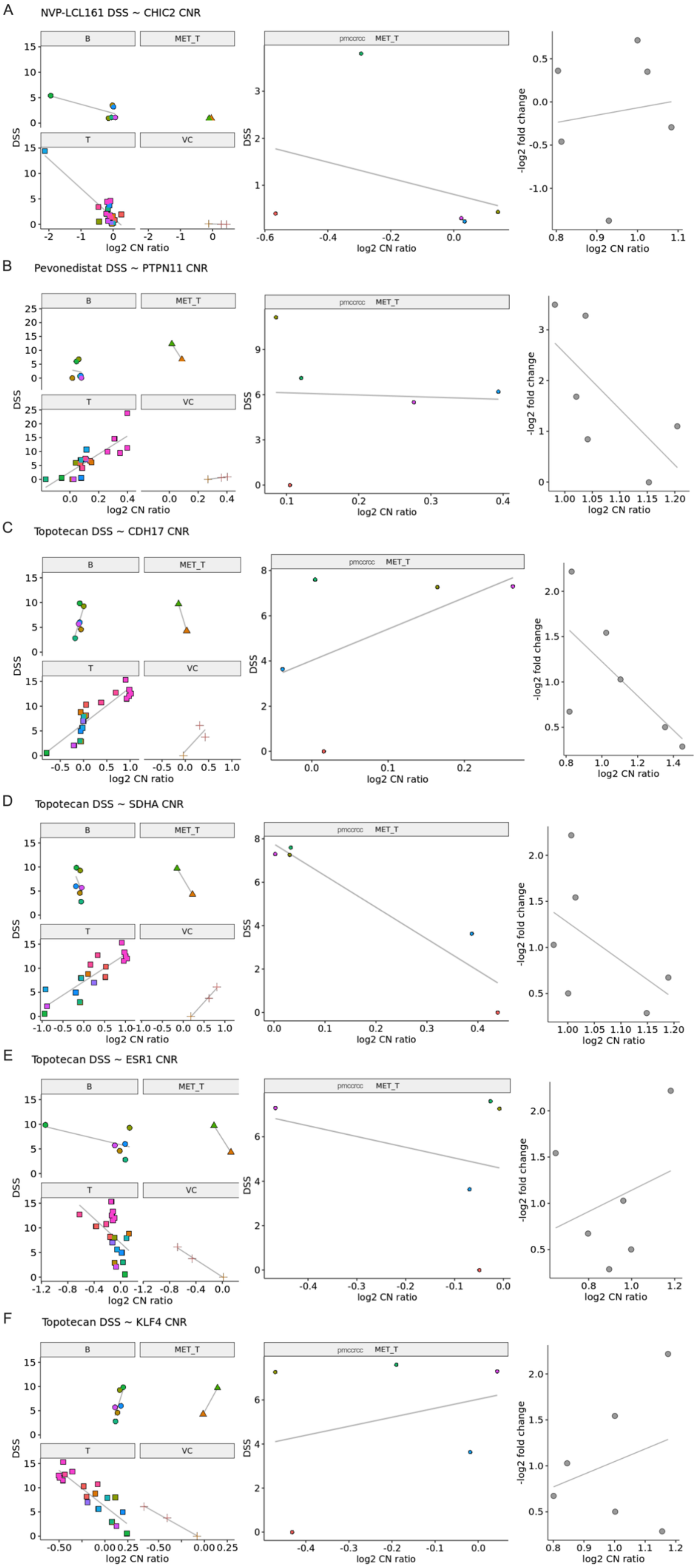
A
NVP-LCL161 DSS ~ CHIC2 CNR
B
Pevonedistat DSS ~ PTPN11 CNR
C
Topotecan DSS ~ CDH17 CNR
D
Topotecan DSS ~ SDHA CNR
E
Topotecan DSS ~ ESR1 CNR
F
Topotecan DSS ~ KLF4 CNR
B
MET_T
T
VC
pmccrcc
DSS
-log2 fold change
log2 CN ratio

***Supplementary Figure 13. Novel genomic biomarker associations with ex vivo drug response in ccRCC PDCs (part 1/3).***

*Ex vivo drug response (DSS or -log2 fold change (-LFC)) ~ CN ratio associations of the following drug/gene pairs:* ***A.*** *NVP-LCL161~CHIC2,* ***B.*** *Pevonedistat~PTPN11,* ***C.*** *Topotecan~CDH17,* ***D.*** *Topotecan~SDHA,* ***E.*** *Topotecan~ESR1, and* ***F.*** *Topotecan~KLF4. Linear associations and trend lines are shown separately for DEDUCER benign (B), primary tumor (T), vena cava (VC) and non-pmccrcc metastatic (MET_T) samples (left-most 2x2 plot of each subfigure), pancreatic metastatic (pmccrcc MET_T) samples (middle plot), and DepMap primary-tumor derived ccRCC cancer cell lines (KMRC1, KMRC3, KMRC20, 786O, OSRC2, 769P; right plot). While T and VC samples tend to follow the same trend, metastatic and cancer cell line samples do not.*

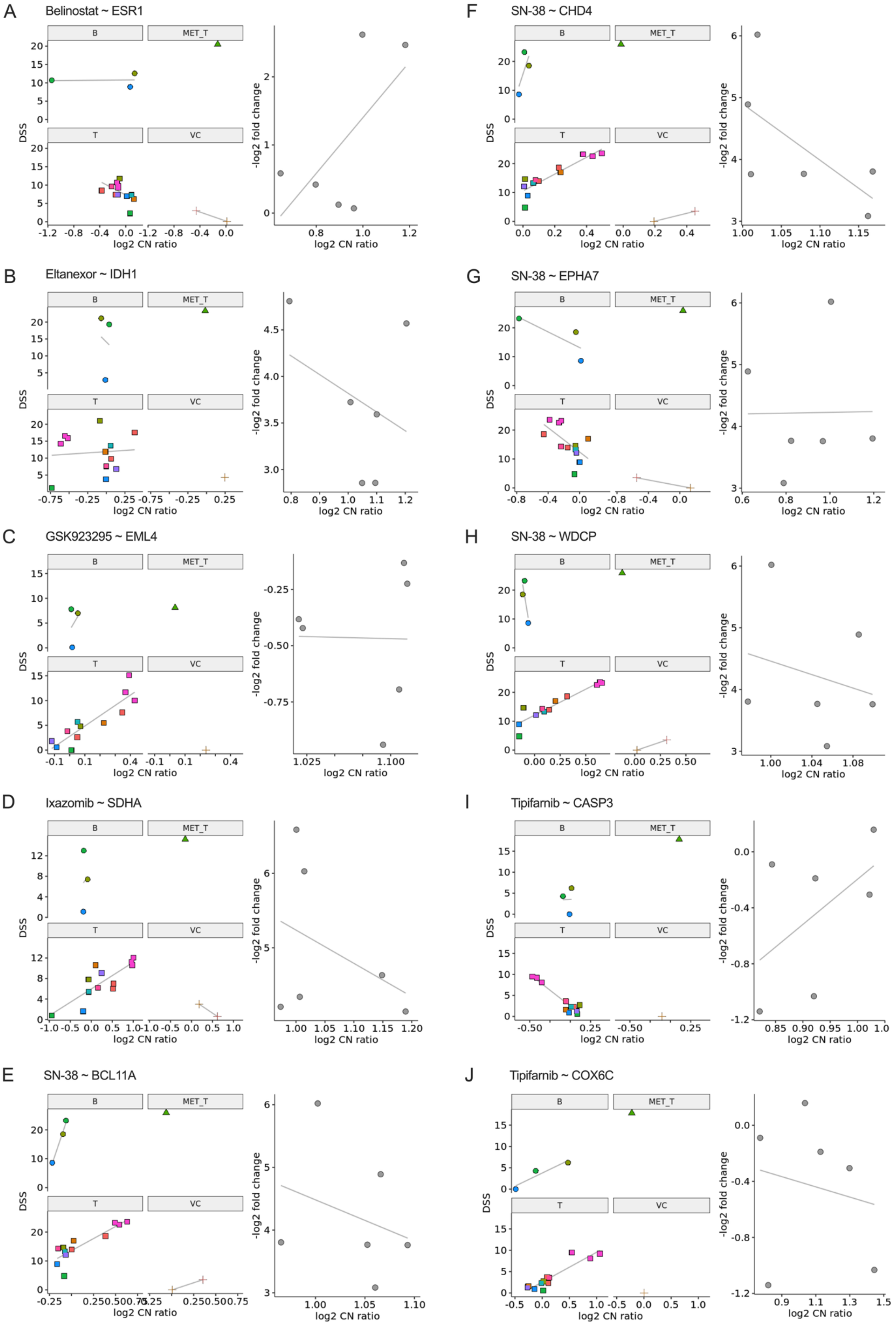

A
Belinostat ~ ESR1
B
Eltanexor ~ IDH1
C
GSK923295 ~ EML4
D
Ixazomib ~ SDHA
E
SN-38 ~ BCL11A
F
SN-38 ~ CHD4
G
SN-38 ~ EPHA7
H
SN-38 ~ WDCP
I
Tipifarnib ~ CASP3
J
Tipifarnib ~ COX6C
B
MET_T
T
VC
DSS
log2 CN ratio
-log2 fold change

***Supplementary Figure 14. Novel genomic biomarker associations with ex vivo drug response in ccRCC PDCs (part 2/3).***

*Ex vivo drug response (DSS or -log2 fold change (-LFC)) ~ CN ratio associations of the following drug/gene pairs:* ***A.*** *Belinostat~ESR1,* ***B.*** *Eltanexor~IDH1,* ***C.*** *GSK923295~EML4,* ***D.*** *Ixazomib~SDHA,* ***E.*** *SN-38~BCL11A,* ***F.*** *SN-38~CHD4,* ***G.*** *SN-38~EPHA7,* ***H.*** *SN-38~WDCP,* ***I.*** *Tipifarnib~CASP3, and* ***J.*** *Tipifarnib~COX6C. Linear associations and trend lines are shown separately for DEDUCER benign (B), primary tumor (T), vena cava (VC) and non-pmccrcc metastatic (MET_T) samples (left-most 2x2 plot of each subfigure), and DepMap primary-tumor derived ccRCC cancer cell lines (KMRC1, KMRC3, KMRC20, 786O, OSRC2, 769P; right plot). While T and VC samples tend to follow the same trend, cancer cell line samples do not.*

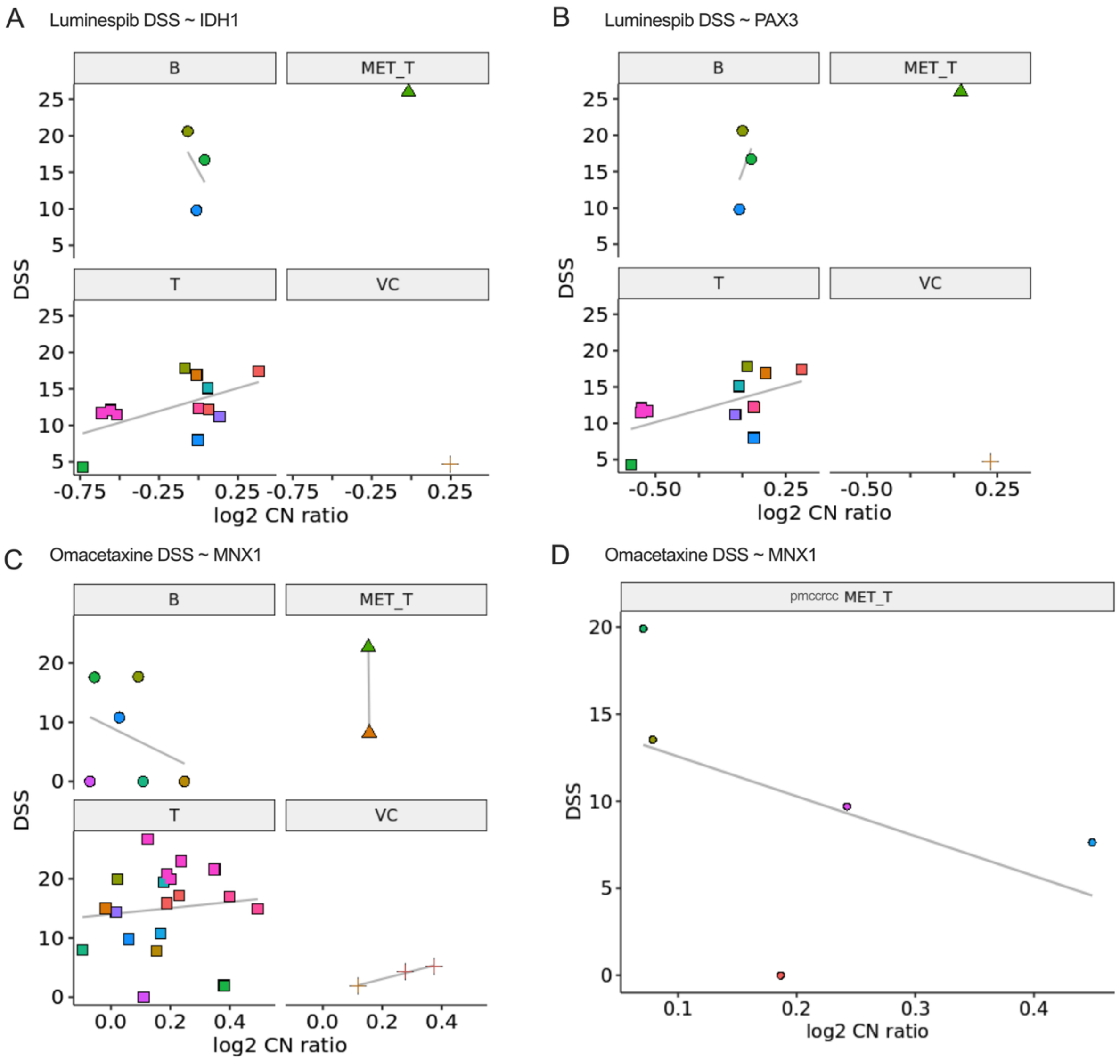


***Supplementary Figure 15. Novel genomic biomarker associations with ex vivo drug response in ccRCC PDCs (part 3/3).***

*Ex vivo drug response (DSS) ~ CN ratio associations of the following drug/gene pairs:* ***A.*** *Luminespib~IDH1,* ***B.*** *Luminespib~PAX3 and* ***C-D.*** *Omacetaxine~MNX1. Linear associations and trend lines are shown separately for DEDUCER benign (B), primary tumor (T), vena cava (VC) and non-pmccrcc metastatic (MET_T) samples (**A-C**), and pancreatic metastatic (pmccrcc MET_T) samples (**D**). While T and VC samples tend to follow the same trend, metastatic samples do not.*

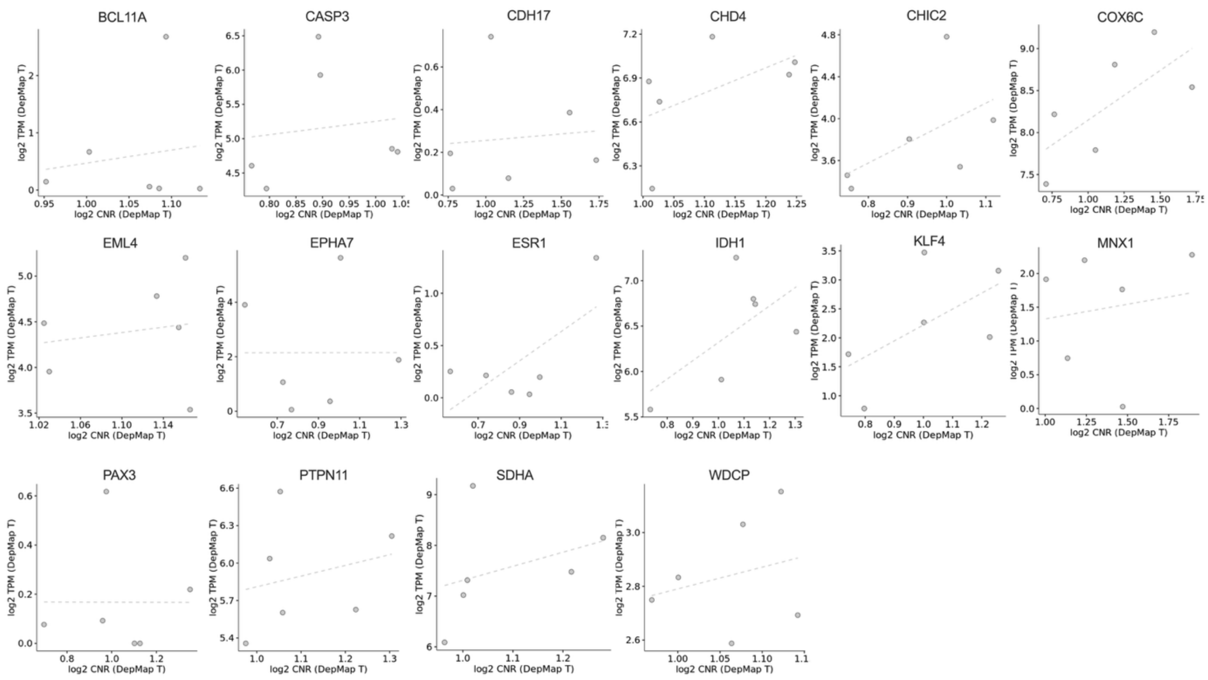


***Supplementary Figure 16. Genomic copy number of novel biomarkers positively correlates with mRNA levels in DepMap ccRCC cancer cell lines.***

*The mRNA expression (TPM) ~ CN ratio associations for 16 biomarker genes with corresponding data available for the DepMap primary-tumor derived ccRCC cancer cell lines (KMRC1, KMRC3, KMRC20, 786O, OSRC2, 769P). In total, 15/16 genes showed a positive association between genomic copy number and transcript expression level.*

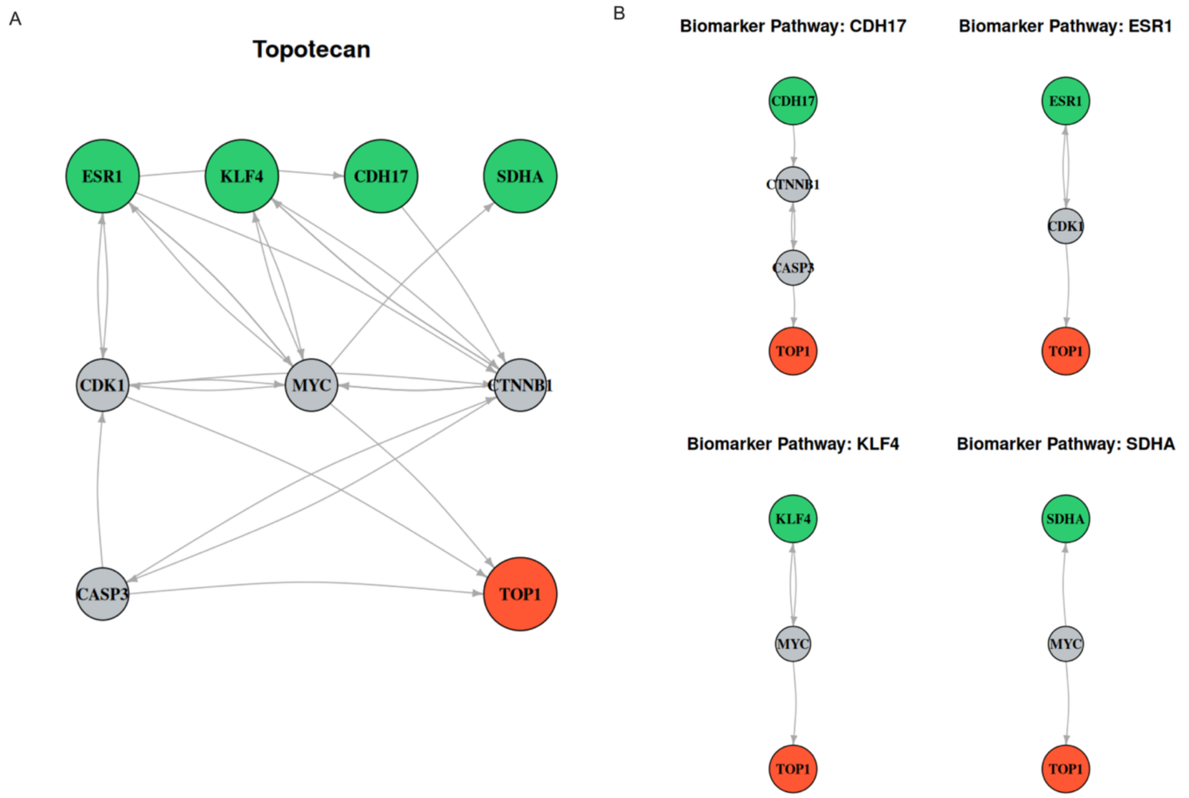


***Supplementary Figure 17. Network connections between the direct Topotecan target and predicted biomarkers.***

*Combined and biomarker-specific shortest-path OmniPath subnetworks linking the direct Topotecan target (TOP1) to the predicted biomarkers (CDH17, ESR1, KLF4, and SDHA). The combined network illustrates distinct signaling routes connecting TOP1 to each biomarker through a small set of shared intermediate nodes, whereas the biomarker-specific subnetworks highlight the corresponding shortest-path connections. Node colors: green, predicted biomarkers; gray, intermediate signaling molecules; red, direct drug target.*
